# Instructive starPEG-Heparin biohybrid 3D cultures for modeling human neural stem cell plasticity, neurogenesis, and neurodegeneration

**DOI:** 10.1101/225243

**Authors:** Christos Papadimitriou, Mehmet I. Cosacak, Violeta Mashkaryan, Hilal Celikkaya, Laura Bray, Prabesh Bhattarai, Heike Hollak, Xin Chen, Shuijin He, Christopher L. Antos, Alvin K. Thomas, Jens Friedrichs, Andreas Dahl, Yixin Zhang, Uwe Freudenberg, Carsten Werner, Caghan Kizil

## Abstract

Three-dimensional models of human neural development and neurodegeneration are crucial when exploring stem-cell-based regenerative therapies in a tissue-mimetic manner. However, existing 3D culture systems are not sufficient to model the inherent plasticity of NSCs due to their ill-defined composition and lack of controllability of the physical properties. Adapting a glycosaminoglycan-based, cell-responsive hydrogel platform, we stimulated primary and induced human neural stem cells (NSCs) to manifest neurogenic plasticity and form extensive neuronal networks *in vitro*. The 3D cultures exhibited neurotransmitter responsiveness, electrophysiological activity, and tissue-specific extracellular matrix (ECM) deposition. By whole transcriptome sequencing, we identified that 3D cultures express mature neuronal markers, and reflect the *in vivo* make-up of mature cortical neurons compared to 2D cultures. Thus, our data suggest that our established 3D hydrogel culture supports the tissue-mimetic maturation of human neurons. We also exemplarily modeled neurodegenerative conditions by treating the cultures with Aβ42 peptide and observed the known human pathological effects of Alzheimer’s disease including reduced NSC proliferation, impaired neuronal network formation, synaptic loss and failure in ECM deposition as well as elevated Tau hyperphosphorylation and formation of neurofibrillary tangles. We determined the changes in transcriptomes of primary and induced NSC-derived neurons after Aβ42, providing a useful resource for further studies. Thus, our hydrogel-based human cortical 3D cell culture is a powerful platform for studying various aspects of neural development and neurodegeneration, as exemplified for Aβ42 toxicity and neurogenic stem cell plasticity.

**Significance:** Neural stem cells (NSC) are reservoir for new neurons in human brains, yet they fail to form neurons after neurodegeneration. Therefore, understanding the potential use of NSCs for stem cell-based regenerative therapies requires tissue-mimetic humanized experimental systems. We report the adaptation of a 3D bio-instructive hydrogel culture system where human NSCs form neurons that later form networks in a controlled microenvironment. We also modeled neurodegenerative toxicity by using Amyloid-beta4 peptide, a hallmark of Alzheimer’s disease, observed phenotypes reminiscent of human brains, and determined the global gene expression changes during development and degeneration of neurons. Thus, our reductionist humanized culture model will be an important tool to address NSC plasticity, neurogenicity, and network formation in health and disease.

## Text

### Introduction

Human brain development and neuronal diseases cannot be modeled adequately by current animal models (LaFerla and Green, 2012); therefore, the development of novel humanized systems that manifest neurogenic plasticity is necessary. The brain’s plasticity provides an endogenous reservoir of cells that could be harnessed to physiologically enhance brain capacity or for neuronal repair (Gage and Temple, 2013; Wyss-Coray, 2016). Therefore, it is fundamentally important to understand how neurons develop to form a hard-wired network, how new networks are generated when newly formed neurons are incorporated, and how stem cells contribute to these processes. Animal models and cell culture experiments examining how mammalian brains develop and elucidating the molecular programs regulating this process have been invaluable (Molyneaux et al., 2007). However, the human brain might exhibit differences compared with the model systems that might not be discernable with existing tools.

The human brain cannot repair the loss of neurons caused by neurodegenerative diseases (ND), in part due to reduced stem cell proliferation and neurogenesis (Tincer et al., 2016). These combinatorial effects exacerbate the manifestation of the disease. In ND states in humans, overall plasticity is severely decreased (Heneka et al., 2015; Nalbantoglu et al., 1997; Selkoe, 2002), however, we know little about how to circumvent this reduction due to lack of appropriate experimental systems. Alzheimer’s disease (AD) is the most common ND, and one of its hallmarks is the aggregation of amyloid protein cleavage products – mainly amyloid-β-42 (Aβ42) peptides (Esler and Wolfe, 2001; Haass and Selkoe, 2007; Nalbantoglu et al., 1997). The disease state produces an inhospitable environment lacking homeostasis in which stem cells are unable to form neurons and new cells do not survive and successfully integrate into existing circuitry. Therefore, understanding stem cell plasticity and neuronal behavior in disease-related settings is critical to determine if a stem cell can regain proliferative and neurogenic function or whether a newborn neuron can survive and integrate into the remaining circuitry despite prevalent amyloid toxicity in the brain.

The overall plasticity of the human brain requires neural stem cell (NSC) proliferation, neurogenesis and neuronal network formation (Alvarez-Buylla et al., 2002; Gage, 2000). However, although NSCs in human brain possess the plasticity to fulfill all these steps, 2D culture conditions are insufficient to generate the connected arbors and long-term behaviors observed in the brain (Haycock, 2011; Justice et al., 2009). Culture conditions that drive the generation of neurons that retain a mature neuronal morphology and form synapses and 3D patterns are needed to address how the entire spectrum of plasticity manifests in the human brain. In addition, paracrine effects from the interplay between glia and neurons, the nascent cellular microenvironment, and the extracellular matrix (ECM) regulate cell fate decisions (Lutolf et al., 2009). In particular, the modulation of mechanical cues, the degradability of the matrix, and the administration of soluble effectors is known to control stem cell fate (Discher et al., 2009). Therefore, we require advanced customized and controllable assay systems for human neural stem cells.

Animal models of AD are unable to recapitulate the entire human disease spectrum (LaFerla and Green, 2012), suggesting that human cells might have a different physiological response than animal cells in response to NDs. Additionally, revolutionary 3D technologies are useful tools for addressing specific questions; however, they require highly complex culture conditions, are difficult to establish and reproduce, and the content of the scaffolds often have unintended consequences on the encapsulated cells. Therefore, *in vitro* systems utilizing human cells to model neural development or neurodegeneration in an *in vivo*-like 3D environment that is amenable to manipulation and monitoring would be highly beneficial. Therefore, in this study, we adapted a highly tunable and defined glycosaminoglycan (GAG)-based 3D matrix system (Chwalek et al., 2014; Maitz et al., 2013) to culture primary and induced human neural stem cells, which manifest their plasticity and neurogenic capacity. This system allowed us to systematically vary the local cellular environment in terms of stiffness, degradability and presentation of GAG-affine signaling molecules (Capilla and Linhardt, 2002) to identify optimal conditions that reproducibly form networks of mature neurons and glia that serve as neural stem cells. We also recapitulated important aspects of AD by modeling amyloid toxicity. Aβ42 treatment impaired network formation and progenitor cell proliferation and induced human pathological hallmarks, including Tau hyperphosphorylation and neurofibrillary tangle formation, suggesting that our biohybrid 3D culture system can be used to address questions regarding neural stem cell proliferation, neurodevelopment and neurodegenerative diseases in a reductionist and tissue mimetic pre-clinical setting.

## Results

To generate a culture system that would allow NSCs to manifest their plasticity and neurogenic capacity in a tissue-representative manner, we applied a modular biohybrid material based on star-shaped poly(ethylene glycol) (starPEG) and GAG heparin (HEP) that can be used to independently tune mechanical cues and biomolecular functionalization (Freudenberg et al., 2012; Tsurkan et al., 2013) when embedding primary human neural stem cells (human cortical astrocytes, from here on primary NSCs) derived from fetal tissue at gestation week 21 (Fig. 1A). We systematically varied the biohybrid matrix in terms of stiffness, cell-responsive remodeling potential, and the presence of soluble effector mediating GAGs to induce the cellular morphology reminiscent of *in vivo* (Supplementary Figure 1). After varying the stiffness (Young’s modulus) of the hydrogels from 0.5 kPa to 3 kPa (Fig. 1B,B’), 1.2 kPa was the optimal stiffness to promote the formation of extended neuronal networks by primary NSCs (Fig. 1B’). Although soft gels (0.5 kPa) disintegrated within 1 week (data not shown), stiff gels (3 kPa) resulted in round nonproliferative cells that do not grow into the matrix or form a network (Fig. 1B’).

**Figure 1.**
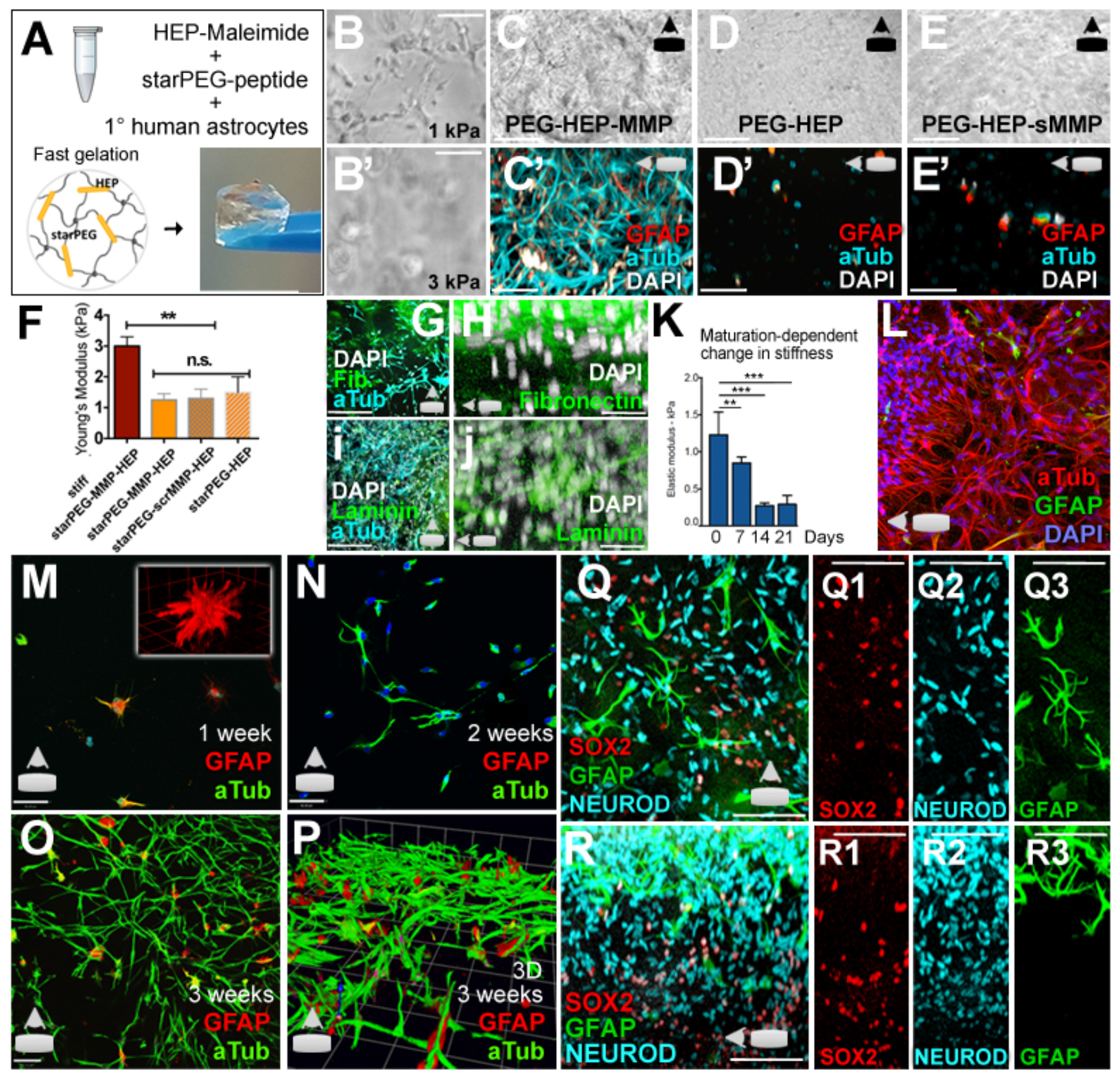
*Physical properties and dynamic remodeling of 3D starPEG-Heparin hydrogels and tissue-mimetic behavior of primary human neural stem/progenitor cells (NSPCs)*. (A) Simplified composition and preparation scheme of glycosaminoglycan (GAG)-based minimalist multifunctional hydrogels. (B, B′) Bright field gel image (BFI) at 1-kPa (B) and 3-kPa (B′) stiffness. (C-E′) BFI (C,D,E) and immunostaining for acetylated-tubulin (aTub) and GFAP (C′,D′,E′) in a 3-week-old gel at 1 kPa stiffness with (C, C′), without (D, D′) and with scrambled MMP cleavage sites (E, E′). (F) Rheological measurements of various gel types. (G) Acetylated-tubulin and fibronectin. (H) Fibronectin. (I) Acetylated-tubulin and laminin. (J) Laminin. (K)Atomic force microscopy-based quantification of elastic modulus in the development of the neural networks. (L) Maximum projection for TUBB3 and GFAP (X-axis). (M-O) Acetylated-tubulin/GFAP staining over the z-axis after 1 (M), 2 (N), and 3 (O) weeks of culture. (P) 3D representation for acetylated-tubulin and GFAP. (Q-R3) Maximum projection images for SOX2, GFAP, and NEUROD1 over the Z-axis (Q-Q3) and X-axis (R-R3). Single fluorescent channels are shown on the right (Q1-Q3, R1-R3). Scale bars: 10 μm for H, J, and M-P; 50 μm for the other figures. Gels: 3 weeks of culture. See also Supplementary Movies 1, 2, and 3.

Tissue characteristics and patterning can be influenced by the cell-responsive remodeling of biohybrid hydrogels, which can be achieved by the incorporation of matrix-metalloprotease-cleavable linkers (Chwalek et al., 2011; Tsurkan et al., 2010). Using MMP-cleavable hydrogels (starPEG-MMP-HEP) (Fig. 1C,C’), directly linked starPEG-HEP gels (Fig. 1D,D’) and non-degradable controls with a scrambled (MMP-insensitive) peptide sequence (starPEG-scr-HEP) (Fig. 1E,E’), we found that only MMP-responsive hydrogels induced the formation of a neuronal network (Fig. 1C). Given that the optimal stiffness was 1200 Pa for all of these gels (Fig. 1F), these results clearly point to enzymatic remodeling involving MMPs (Agrawal et al., 2008) as a crucial process for axodendritic outgrowth and the formation of mature arbors. Moreover, if an enzymatically degradable hydrogel of inert starPEG was formed, the cells did not proliferate and did not display axodendritic extensions, highlighting the importance of a factor regulating GAG-heparin interaction within the matrices (data not shown). Because heparin can bind multiple insoluble matrix proteins (e.g., laminin, fibronectin) and other soluble growth factors (Capila and Linhardt, 2002; Garg et al., 2011), it might mediate the activity and presentation of cell-secreted molecules and thus indirectly control cell fate processes. Sulfated GAGs, particularly heparan sulfate and chondroitin sulfate, are well known to be a major component of neuronal ECMs and are crucial for developmental processes and axon guidance (Lau et al., 2013). Thus, the MMP-cleavable heparin hydrogel matrix provided optimal conditions with respect to initial stiffness and the presence of sulfated GAGs and promoted remodeling in a well-orchestrated and timely way via MMP-sensitive peptide linkers.

Since the development of 3D cultures requires MMP-cleavage, we hypothesized that the observed maturation and patterning might follow a replacement of the initial scaffold with the cells’ own matrix. Therefore, we immunostained the cultures for fibronectin and laminin to test this hypothesis and found that the 3D gels generate fibronectin (Fig. 1G,H) and laminin (Fig. 1I,J) *de novo*, suggesting that neural stem cells and neurons remodel the matrix allowing to generate a liberal stem cell niche according to the needs of the NSCs. To investigate whether this remodeling would alter the stiffness of the gel matrix, we performed atomic force microscopy analyses of the gels at different time points and observed that the initial elastic modulus of 1.3 kPa is reduced gradually to 0.3 kPa after 14 days (Fig. 1K), indicating that the development of starPEG-Heparin 3D cultures generate human brain-mimetic physical properties. This is particularly important because widely used 3D scaffolds (e.g.: Matrigel) cannot be regulated in their stiffness by the encapsulated cells, and therefore they deviate from tissue-mimetic physical features. Indeed, when we compared the network forming ability of primary human NSCs in identical culture conditions in starPEG-Heparin and Matrigel, we found that the neurogenic capacity and network-formation ability of primary NSCs manifest significantly better in starPEG-Heparin gels (Supplementary Figure 2), indicating that starPEG-Heparin composition favors an unprecedented tissue-mimetic environment for primary NSCs.

Based on our cell-matrix interaction results, we hypothesized that the maturation of neuronal networks (Fig. 1L) might resemble human neurodevelopment. We analyzed the gels at various time points to address this hypothesis (Fig. 1M–P). One week after seeding, the gel contained sparsely distributed GFAP-positive glia with a 3D arborized morphology (Fig. 1M). After two weeks of culture, we started to observe acetylated tubulin-positive neurons extending processes and organizing themselves into clusters (Fig. 1N; Suppl. Video 1). At 3 weeks, the cultures produced an extensive and elaborate network of neurons with interspersed glia (Fig. 1O) in close association with neurons (Fig. 1P). We also determined that the primary NSC cultures expressed neural stem cell markers such as SOX2 and GFAP, and neural fate determinants such as NEUROD in spatially overlapping, but distinct domains (Fig. 1Q, X-axis view; Fig. 1R, Z-axis view), indicating that neural stem cell plasticity programs and neurogenic activity drive the development of the neuronal networks, and the stem cell compartments may pattern in a 3D topology similar to human brains.

Based on our findings, we hypothesized that 3D cultures would provide a superior 3D topological environment to pNSCs, and therefore would instruct a gene expression profile that would be closer to in vivo. To investigate how 3D cultures would molecularly differ from 2D cultures, we cultured primary NSCs in 2D and 3D using identical culture conditions for three weeks. After isolating total mRNA, we performed whole transcriptome sequencing, and observed that a considerable number of genes are differentially expressed between 2D and 3D cultures (Fig. 2A, Supplementary Dataset 1). In order to identify the pathways and molecular programs represented better in 3D cultures, we performed pathway and enrichment analyses (Fig. 2B). We found that 3D cultures express genes related to several pathways that characterize mature neuronal physiology such as focal adhesion, ECM-receptor interaction, axon guidance, and various signaling pathways (Fig. 2B, Suppl. Dataset 2). Additionally, cellular component analyses indicate that the 3D cultures express the genes, the protein products of which are related to various mature neuronal processes such as synapses and axons (Fig. 2B). These results indicate that our hydrogel cultures provide the 3D topology and instructive environment to generate neuronal networks from primary human NSCs in a tissue-mimetic manner, which is not the case for 2D cultures.

**Figure 2.**
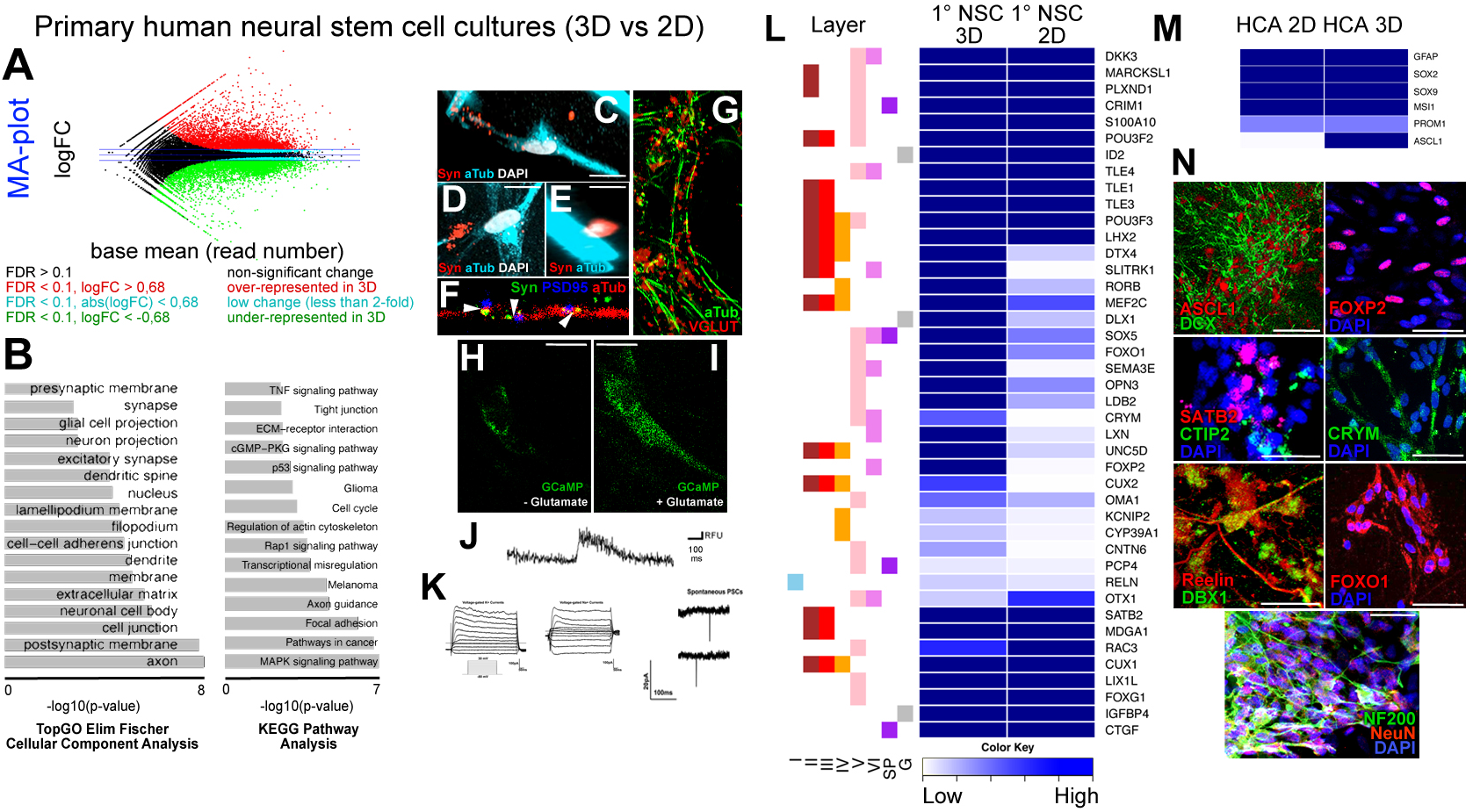
*Comparison of gene expression profiles of primary human cortical astrocytes in 2D and 3D cultures*. (A) MA-plot for differentially expressed genes. Red: upregulated in 3D, green: downregulated in 3D. (B) KEGG pathway analyses pie chart showing significantly enriched molecular pathways. (C-E) Synaptophysin and Acetylated tubulin immunostaining in 3D cultures. (F) Synaptophysin, PSD95 and Acetylated tubulin immunostaining in 3D cultures. (G) VGLUT1 and Acetylated tubulin immunostaining in 3D cultures. (H) GCaMP6 signal without Glutamate treatment. (I) GCaMP6 signal with Glutamate treatment. (J) Fluorescence intensity histogram. Note the peak at the time of Glutamate treatment. (K) Patch clamp recordings for Na+ (left) and K+ (middle) currents. Detection of spontaneous firing of neurons (right). (L) Heat map for expression levels of cortical marker genes for 2D and 3D cultures. Asterisks indicate significant difference between two samples. Genes are denoted with their respective cortical layer expression with color codes (left to the heat map). (M) Heat map for expression levels of neural stem cell markers. (N) Immunostaining for ASCL1 (proneural determinant), DCX (early neuronal marker), SATB2, CTIP2, Reelin (RELN), DBX1, FOXP2, CRYM, FOXO1 (cortical markers), and Neurofilament (NF200) and NeuN (mature neuronal markers). Scale bars: 10 μm for C-I; 25 μm for N. Gels: 3 weeks of culture. See also Supplementary Datasets 1, 2 and 3.

To verify that the 3D cultures of primary human NSCs generate mature neurons resembling the *in vivo* conditions, we immunostained 3D cultures for the synaptic marker Synaptophysin (Fig. 2C), which clusters at neuronal junctions (Fig. 2D) and boutons (Fig. 2E), and generate pre- and post-synaptic termini (Fig. 2F), indicating that neurons in 3D cultures develop enough to form synaptic connections, which we did not observe in 2D conditions (data not shown). We found that the neurons formed in the 3D starPEG-HEP-based hydrogels were also expressing neurotransmitter receptors such as VGLUT1 (Fig. 2G), and are responsive to neurotransmitters such as glutamate, as shown by increased intracellular calcium levels (Fig. 2H–J; Suppl. Video 2) following the transfection of plasmids expressing the GCamP6f calcium sensor driven by the CMV promoter. We also performed electrophysiology experiments assessing sodium and potassium channel activity to determine whether the cells from the matrices were functional neurons. We observed potassium and sodium channel activity as well as spontaneous firing of neurons in whole cell patch clamp recordings (Fig. 2K), indicating that cells cultured within the hydrogels differentiated into functionally active neurons. These results indicate that starPEG-HEP-hydrogel-based 3D cultures of primary human NSCs are able to generate an elaborate and mature network of neurons and glia in a three dimensional organization.

To determine the cortical subtypes produced in 3D and 2D conditions, we compared the expression levels of a selected set of cortical marker genes in relation to their spatial confinement to different layers (Fig. 2L; from (Molyneaux et al., 2007)). We observed that 3D conditions favor for expression of a larger and more complete set of cortical marker genes especially associated with layers IV, V and VI, compared to 2D (Fig. 2L). Similarly, although 2D and 3D conditions allow expression of NSC markers at high levels, the major cortical pro-neural gene ASCL1 is expressed in rather low levels in 2D conditions while 3D cultures allow significantly higher levels of ASCL1 expression (Fig. 2M). To validate our deep sequencing results and heat map analyses, we performed immunohistochemistry for pro-neural determinant ASCL1 and early neuronal marker DCX, cortical layer markers SATB2, CTIP2, Reelin (RELN), DBX1, FOXO1, FOXP2, CRYM, and mature neuronal markers Neurofilament-200 and NeuN (Fig. 2N). We found that 3D cultures allow expression of pro-neural fate determinants, and production of different lineage subtypes of cortical neurons (Fig. 2N).

To test whether we could use our system with other types of neural stem cells, we cultured iPSC-derived human neural stem cells (iNSCs) in culture under conditions identical to those of primary NSCs (pNSCs), and observed extensive neurogenesis and network formation similar to pNSCs at 3 weeks of cultures (Fig. 3A,B, Suppl. Movie 3), suggesting that our instructive hydrogel composition can support development of networks from various types of neural stem cell sources.

**Figure 3.**
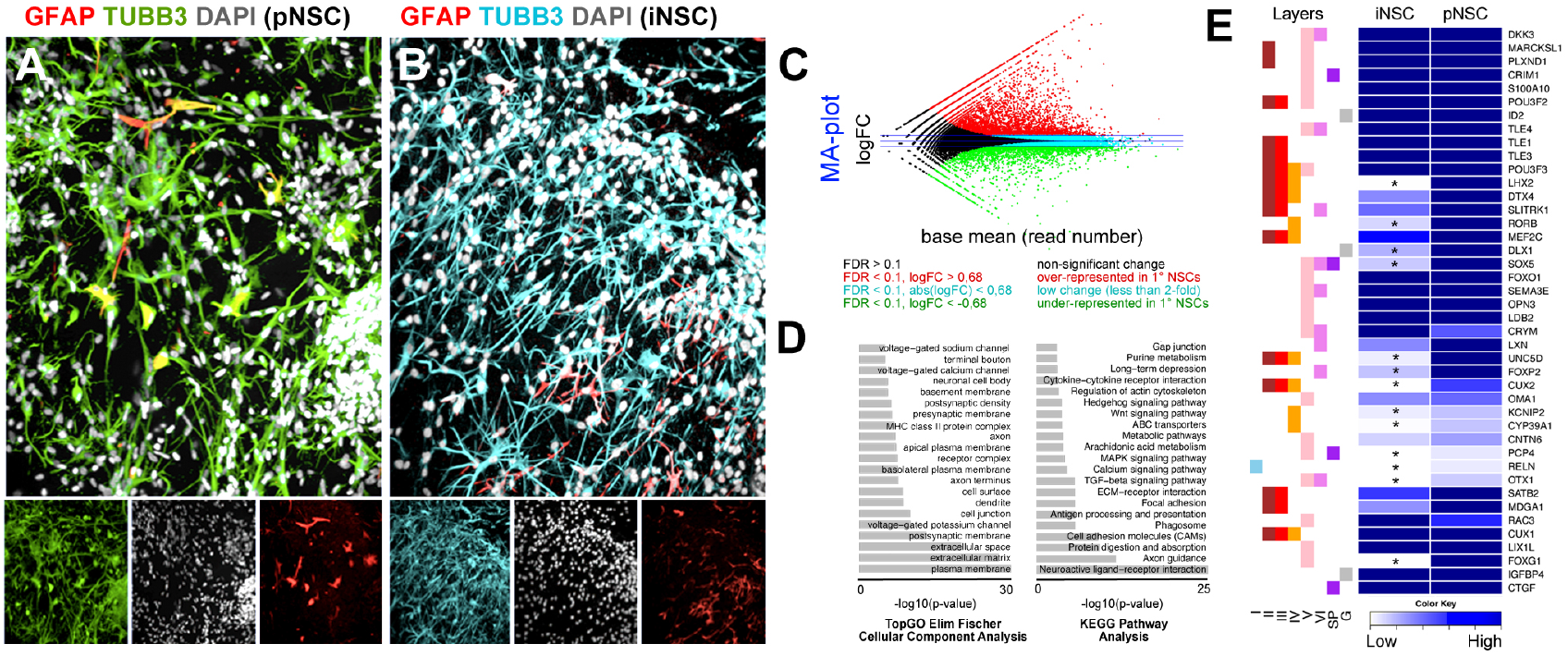
*Culture of iPSC-derived neural stem cells in 3D, and comparison of whole transcriptome profile to primary human neural stem cell-derived cultures*. (A) Immunostaining for GFAP (red) and TUBB3 (green) on 3 weeks of 3D cultures with primary human cortical astrocytes (neural stem cells). Insets below the panel are individual fluorescence channels including DAPI (white). (B) Immunostaining for GFAP (red) and TUBB3 (cyan) on 3 weeks of 3D cultures with iPSC-derived human neural stem cells. Insets below the panel are individual fluorescence channels including DAPI (white). (C) MA-plot for differentially expressed genes in primary over iPSC-derived NSC cultures. Red: over-represented in primary cultures, green: underrepresented in primary cultures compared to iPSC cultures. (D) KEGG pathway analyses pie chart showing significantly enriched molecular pathways in primary cultures compared to iPSC cultures. (E) Heat map for expression levels of cortical marker genes in iPSC and primary cultures. Asterisks indicate significant difference between two samples. Genes are denoted with their respective cortical layer expression with color codes (left to the heat map). Scale bars: 100 μm. Gels: 3 weeks of culture. See also Supplementary Datasets 4, 5 and 6.

To compare the molecular expression profiles of pNSC and iNSC cultures, we isolated total RNA and performed whole transcriptome sequencing on iNSCs and compared the reads to those of the cultures with pNSCs (Fig. 3C, Suppl. Dataset 3). We found that although both iNSCs and pNSCs can generate neuronal networks in 3D cultures, these two cell types differ significantly in their gene expression patterns (Fig. 3C; Suppl. Dataset 4). Interestingly, we found that compared to iNSCs, pNSC cultures express genes related to extracellular matrix and plasma membrane more (Fig. 3D; Suppl. Dataset 5), and these differences enrich pathways such as various signaling pathways, axon guidance, neuroactive ligand-receptor interaction and metabolism (Fig. 3D; Suppl. Dataset 6). Furthermore, when we compared cortical layer marker expression in iNSCs and pNSCs, we found that compared to pNSCs, iNSC cultures cannot form a subset of neuronal lineages especially for layers II, III and IV in our particular culture conditions (Fig. 3E). These findings suggest that our 3D culture system can be used to dissect the properties and neurogenic capacities of different progenitor types in particular culture settings and under certain physical parameters, and may serve as a suitable tool for investigating the physiological differences between induced and primary NSC populations.

Since our culture system can form cortical neurons, the development of which relies on the plasticity and neurogenic ability of human NSCs, we hypothesized that we could model disease conditions that lead to impaired NSC plasticity and neurogenic output. In the human brain, Aβ42 aggregation impairs neuronal network formation and neuronal connectivity due to death of existing neurons as well as loss of neurogenic ability and neural stem cell plasticity (Hardy and Selkoe, 2002; Kienlen-Campard et al., 2002; LaFerla et al., 2007; Selkoe, 2002; Tincer et al., 2016). Aβ42 was shown to negatively affect the plasticity and neurogenesis of NSCs in mouse models of AD (Ermini et al., 2008; Haughey et al., 2002; Heo et al., 2007), and the same effect is one of the principle limitations of current neuro-regenerative approaches to the treatment of AD in humans (Cosacak et al., 2015; Demars et al., 2010; Tincer et al., 2016). However, the mechanisms underlying the impact of Aβ42 on NSC plasticity are still largely unknown and cannot be elucidated analytically in human brains. We therefore extended our above-described 3D cultures by treating primary NSCs with Aβ42, a major hallmark of AD pathology, before embedding them in the biohybrid hydrogels (Figure 4A). We used an Aβ42 form that we previously found to be causing pathological outcomes in vertebrate brains (Bhattarai et al., 2016; Bhattarai et al., 2017b). In control gels, the neurons formed highly connected networks (Fig. 4B) that were disrupted upon Aβ42 treatment (Fig. 4C). We developed an algorithm to trace the connected neuronal paths as skeletonized arbors and quantified the extent of the neuronal connections, and observed that neuronal networks reduce significantly after Amyloid toxicity (Fig. 4D–F). Similar to human brains, Aβ42 resulted in dystrophic axons (Fig. 4G–G’’; Supplementary Video 5), impaired ECM composition and stiffness (Figure 4H–J), Tau hyperphosphorylation (Figure 4K–N, Supplementary Movie 5), neurofibrillary tangle formation as observed by Gallyas silver impregnation (Fig. 4m, Suppl. Fig. 3) and Thioflavin S staining (Fig. 4N), microtubule disassembly (Figure 4O), and amyloid aggregation and autophagy (Figure 4P,Q).

**Figure 4.**
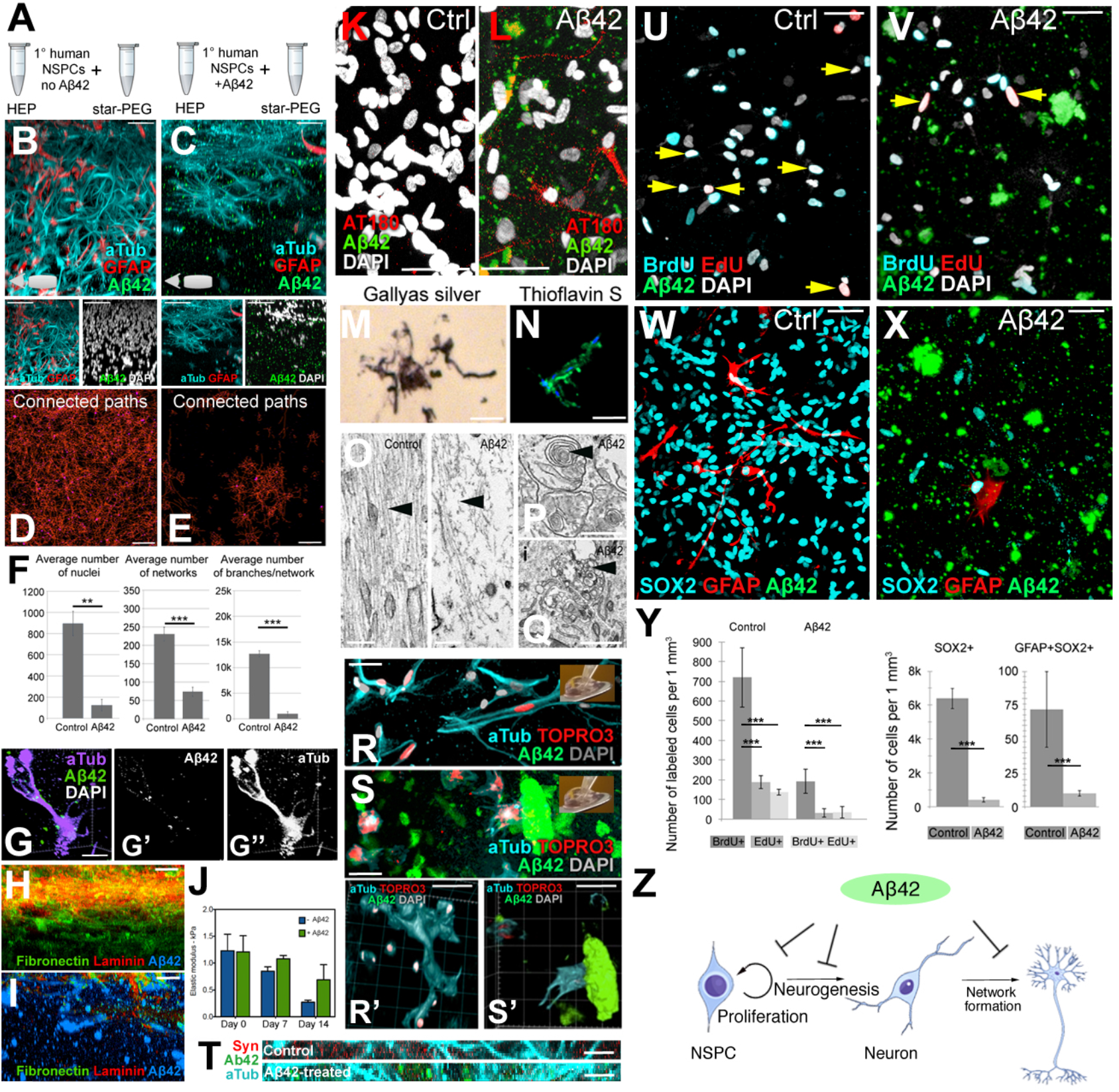
*3D hydrogel cultures of primary human NSCs as an Aβ42 toxicity model*. (A) Simplified gel preparation/Aβ42 administration scheme. (B,C) X-axis views for Aβ42, Acetylated-tubulin, GFAP in control (B) and Aβ42 gels (C). Double stainings under the panels. (D,E) Maximum projection of the skeletonized-connected neuronal pathways in control (D) and Aβ42 gels (E). (F) Quantification. (G-G’’) 3D-reconstruction of TUBB3/Aβ42 showing dystrophic axons. (H,I) Fibronectin/laminin in control (H) and Aβ42 gels (I). (J) Time-course atomic force microscopy measurements of stiffness. (K,L) Phospho-Tau (AT180) in control (K) and Aβ42 gels (L). (M) Gallyas silver staining. (N) Thioflavin-S staining in Aβ42 gels. (O) EM images of microtubules in control (left) and Aβ42 gels (right). (P,Q) Aβ42 depositions (P), and autophagic vacuoles (Q). (R-S’) Acetylated-tubulin and Aβ42 in control (R) and Aβ42 gels (S) after transplantation. Transplanted cell nuclei labeled with TOPRO3 (red). 3D-reconstruction of R (R’) and S (S’). (T) Acetylated-tubulin, synaptophysin, Aβ42 in control (upper) and Aβ42 gels (lower). (U,V) BrdU and EdU in control (U) and Aβ42 gels (V). Arrows: double-positive cells. (W,X) SOX2, GFAP, Aβ42 in control (W) and Aβ42 gels (X). (Y) Quantification of U-X. (Z) Schematics for effects of Aβ42. Scale bars: 10 μm for G, M, N, T, R, and S; 200 nm for O-Q; 50 μm for the other figures. Gels: 3 weeks of culture. See also Supplementary Movies 4-8.

Amyloid toxicity not only impairs neurogenesis and neuronal survival but also reduces synaptic plasticity (Selkoe, 2002). Amyloid load prevents the formation of new synapses, and newly added cells cannot integrate into the circuitry, rendering exogenous stem cell therapy inefficient (Lilja et al., 2015; Tong et al., 2015). We developed a transplantation paradigm with cultured gels to test whether our 3D culture model could be used to address questions regarding the neurogenic potential and capacity of transplanted cells to integrate into existing networks (Fig. 4R). We labeled all pNSCs with a nuclear stain, and injected them into another hydrogel that had been pre-cultured with embedded pNSCs for 1 week (Fig. 4R). At 1 week after transplantation, the injected cells formed neurons with arbors (red nuclei, Fig. 4R; Supplementary Video 7) and connected to preexisting cells in the control gels (Fig. 4R,R’). In contrast, cells injected into Aβ42-containing gels did not acquire an arborized morphology or connect to the existing cells that were also not arborized (Fig. 4S,S’; Supplementary Video 8). Combined with the findings that Aβ42 impairs the synaptic connections overall (Fig. 4T), these results suggest that 3D cultures can be used for analyzing how synaptic connections can be regenerated and how new neurons can be forced to integrate into the existing circuitry upon Aβ42 toxicity.

Aβ42 reduces the NSC plasticity and neurogenic capacity in human brains, and stem cell-based regenerative therapies would require therapeutic activation of NSCs (Tincer et al., 2016). To investigate whether Aβ42 reduced NSC plasticity in our 3D cultures, we determined the proliferative capacity and prevalence of pNSCs after Aβ42 by BrdU/EdU treatment and immunohistochemical stainings for NSC markers SOX2 and GFAP (Fig. 4U–X). We treated the cultures with BrdU at 1 week of development and with EdU at 2 weeks of development, and analyzed the presence of BrdU and EdU positive cells at 3 weeks of cultures where double-positive cells would indicate constitutively proliferating stem cells (Fig. 4U,V). We found that Aβ42 reduced BrdU-EdU incorporation and the number of constitutively proliferating cells (Fig. 4Y) as well as reducing numbers of GFAP and SOX2-positive NSCs (Fig. 4Y), indicating that our 3D culture system can be used for modeling Aβ42-induced impairment of NSC plasticity. Thus, our 3D cultures of NSCs can also successfully serve as a novel *in vitro* sporadic AD model of Aβ42 toxicity on human NSC plasticity and neurogenesis (Figure 4Z). Importantly, we found that all those effects of Aβ42 observed in 3D cultures are specific because scrambled Aβ42 or other Aβ species such as Aβ38 do not show any phenotypes above (Suppl. Figure 4).

To determine if Aβ42 would affect the neuronal network formation and NSC plasticity in iPSC-derived cultures, we treated the iNSCs with Aβ42, encapsulated in 3D gels. To determine the neuronal network formation, NSC prevalence, and proliferative capacity, we performed immmunostainings for GFAP and TUBB3 (Fig. 5A,B), SOX2 and GFAP (Fig. 5C,D), and GFAP and BrdU (Fig. 5E,F) together with Aβ42 detection. Compared to controls, Aβ42 reduces the total number of newborn cells and NSCs (GFAP/SOX2-positive) (Fig. 5G), average number of networks (Fig. 5H), average number of branches per network (Fig. 5I), average branch length per network (Fig. 5J), length of longest connected path (Fig. 5K), and maximum branch length (Fig. 5L) in cultures started with iNSCs. These results show that Aβ42 impairs the plasticity, neurogenic ability and network forming capacity of primary and human NSC cultures in our 3D hydrogel matrix, and our system can also be used for iPSC-derived cell types.

**Figure 5.**
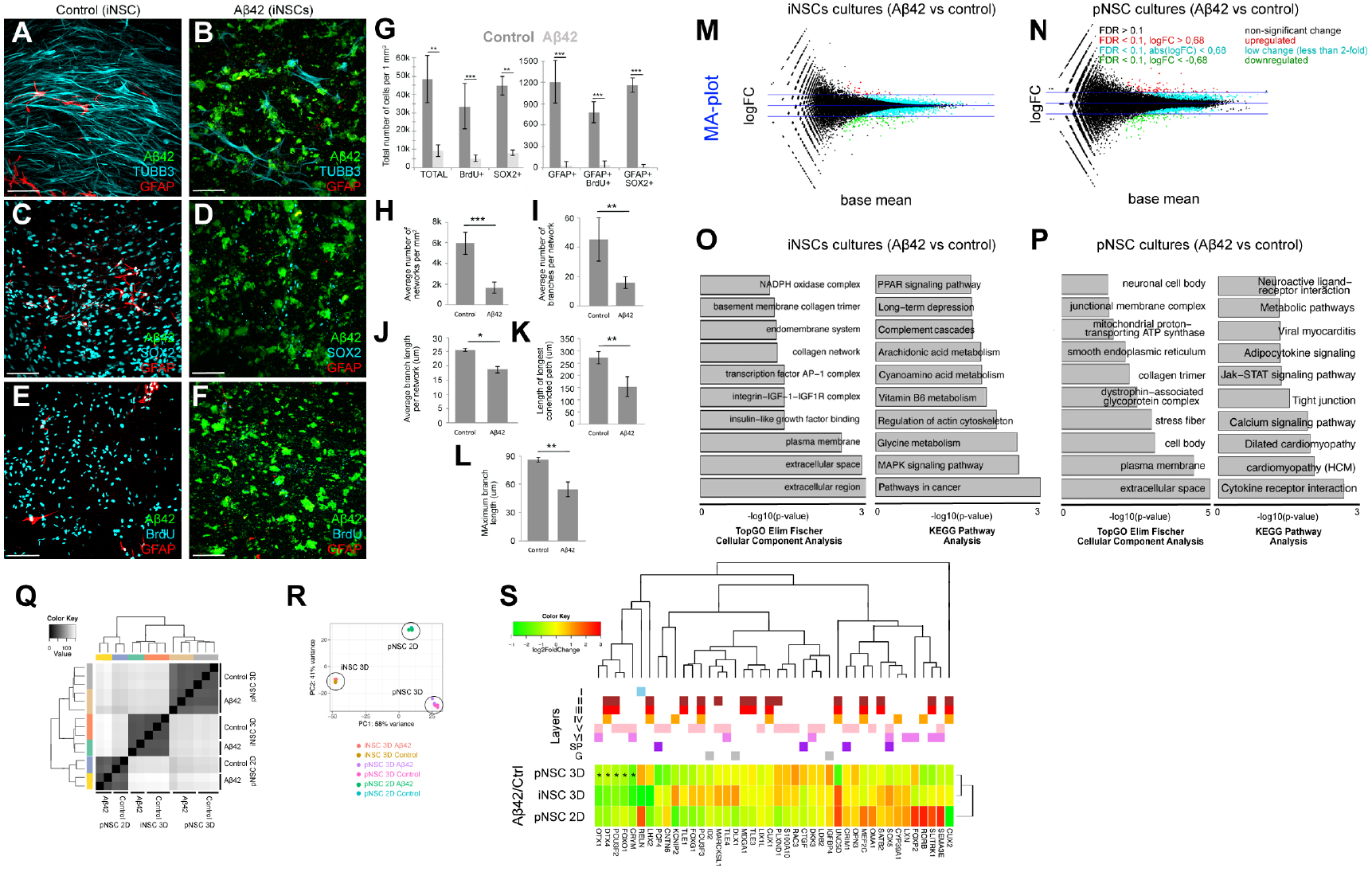
Aβ42 toxicity model with iPSC-derived neural stem cells in 3D hydrogels, analysis of transcriptional changes, and comparison to pNSC cultures. (A) Immunostaining for Aβ42 (green), TUBB3 (cyan) and GFAP (red) on control iPSC-derived NSC cultures. (B) Immunostaining for Aβ42 (green), TUBB3 (cyan) and GFAP (red) on Aβ42-treated iPSC-derived NSC cultures. (C) Immunostaining for Aβ42 (green), SOX2 (cyan) and GFAP (red) on control iPSC-derived NSC cultures. (D) Immunostaining for Aβ42 (green), SOX2 (cyan) and GFAP (red) on Aβ42-treated iPSC-derived NSC cultures. (E) Immunostaining for Aβ42 (green), BrdU (cyan) and GFAP (red) on control iPSC-derived NSC cultures. BrdU is given at 1 week of culture. (F) Immunostaining for Aβ42 (green), TUBB3 (cyan) and GFAP (red) on Aβ42-treated iPSC-derived NSC cultures. BrdU is given at 1 week of culture. (G) Quantification graph for number of cells in control and Aβ42-treated cultures. (H-L) Quantification of average number of networks (H), average number of branches per network (I), average branch length per network (J), length of longest connected path (K), and maximum branch length (L). (M) MA-plot for differentially expressed genes in iPSC-derived NSC cultures after Aβ42 treatment. Red: upregulated, green: downregulated. (N) MA-plot for differentially expressed genes in primary NSC cultures after Aβ42 treatment. Red: upregulated, green: downregulated. (O) KEGG pathway analyses pie chart showing significantly enriched molecular pathways in iPSC cultures after Aβ42 treatment.. (P) KEGG pathway analyses pie chart showing significantly enriched molecular pathways in primary cultures after Aβ42 treatment. (Q) Heat map for changes in expression levels of cortical marker genes in iPSC and primary cultures after Aβ42 treatment. Genes are denoted with their respective cortical layer expression with color codes (above the heat map). Scale bars: 100 μm. Gels: 3 weeks of culture. See also Supplementary Datasets 7-12.

To determine the gene expression changes exerted by Aβ42 in 3D cultures, we performed whole transcriptome sequencing on control and Aβ42-treated cultures initiated with human iNSCs (Fig. 5M, Suppl. Dataset 7) and pNSCs (Fig. 5N, Suppl. Dataset 8). Cellular component analyses and KEGG pathway enrichment analyses in iNSC-derived (Fig. 5O, Suppl. Dataset 9, 10) and primary (Fig. 5P, Suppl. Dataset 11, 12) cultures showed that divergent pathways are affected by Aβ42 in 3D cultures. Hierarchical clustering (Fig. 5Q) and multivariance analyses (Fig. 5R) indicated that primary and induced NSC cultures have their own molecular signatures of gene expression, which are affected by Aβ42. By plotting a heat map of gene expression changes, we also found that several cortical marker genes are differentially expressed after Aβ42 in induced and primary human NSC-based 3D cultures (Fig. 5S), which is suggestive of the alterations in the cortical neuronal subtypes (for instance, POU3F2, CRYM, and FOXO1). Our results indicate that although Aβ42 causes impaired neural stem cell plasticity, neurogenesis and network formation in both iNSC- and pNSC-derived cultures (Fig. 4, Fig. 5), the molecular programs it alters in these cultures do differ (Fig. 5O, P, S). This finding is consistent with previous documentations that primary and iPSC-derived neural stem cells have profound physiological differences that might affect subsequent global gene expression, neuronal maturation capacity, and resilience to disease conditions (Kim et al., 2011; Kim et al., 2010; Verpelli et al., 2013; Xia et al., 2016). Therefore, our results suggest that starPEG-Heparin 3D culture system can also be used to dissect the effects of Aβ42 on different stem cell and neuronal populations. Furthermore, our 3D cultures may help refining the overall toxicity of Aβ42 to distinct physiological states and cellular characteristics of experimental cellular systems. Since our cultures do not contain inflammatory cells, this system will also help to investigate the direct roles of Aβ42 on stem cells and neurons in a reductionist and dissective manner.

## Discussion

Neurodegenerative diseases such as AD present with a perplexing set of impairments, including neuronal death, synaptic degeneration, and the inability of stem cells to produce neurons to replace lost neurons (Kienlen-Campard et al., 2002; LaFerla et al., 2007; Lindvall and Kokaia, 2006; Selkoe, 2002, 2003). Thus, designing effective regenerative therapies for patients with AD requires assay systems that address the parameters of neurodegenerative pathology individually and in combination in a reductionist way using human cells. Rodent models of neurodegeneration cannot recapitulate various aspects of human pathology (Gotz and Ittner, 2008; LaFerla and Green, 2012). Thus, we used primary and iPSC-derived human neural stem cells (iNSCs) in 3D cultures to better reflect neurodevelopmental paradigms and neurodegenerative processes in a reductionist manner. 3D cultures using a well-defined biohybrid hydrogel system based on star-shaped PEG and heparin allowed the generation of extensive neuronal networks that could be used to address neurogenesis and connectivity-based questions.

Existing 3D culture systems, including Matrigel^™^-based cultures, are chemically undefined and heterogeneous in composition and cannot be modified for various parameters, such as stiffness, scaffold composition or bioresponsiveness (Ravi et al., 2015). Thus, the interpretation of often quite variable results is difficult, and it is rarely possible to dissect the influences of different exogenous and paracrine signals on cellular development in an isolated and controllable experimental setup. Our biohybrid hydrogel system based on heparin and star-shaped PEG provides valuable advantages by enabling the independent and ongoing adjustment of biophysical and biomolecular matrix signals. Indeed, we determined that starPEG-Heparin gels allow faster and more elaborate network formation compared to Matrigel upon identical culture conditions (Suppl. Fig 2). Moreover, the method we use to generate the gels does not cause cell death or DNA mutation that occurs in other matrices through the formation of free radicals upon polymer network formation.

Various 3D systems, including organoids, cannot form reproducibly sized or formed structures (Fatehullah et al., 2016). However, MMP-sensitive, starPEG-heparin-based 3D cultures can be customized for all of these parameters, providing better-defined conditions. Compared to previous reports modeling AD in 3D cultures using Matrigel (Choi et al., 2014; Smith et al., 2015), despite differences in the initial cell source, starPEG-heparin 3D gels enabled significantly faster development of neuronal networks that offer various advantages such as high-throughput screening approaches that are suitable for drug discovery.

The composition and architecture of the ECM is an integral parameter governing stem cell activity and tissue modeling. However, due to the complex interplay between multiple ECM-derived signals and their pleiotropic effects, *in vivo* assays pose a challenge to identifying the roles of exogenous cues in tissue patterning. Previously described methods for the formation of three-dimensional neuronal networks (Choi et al., 2014; Fatehullah et al., 2016; Tang-Schomer et al., 2014; Zhang et al., 2014) lack a defined composition and the ability to tune the artificial extracellular matrix. Our novel 3D matrix platform would be advantageous to dissect the roles of matrix properties in stem cell activity and differentiation, because cells can dynamically interact with the scaffold to generate their “own” cell-secreted ECM. Additionally, our hydrogels can be covalently functionalized with different matrix-derived peptides or could be used for the effective administration of GAG-affine soluble signal molecules. By doing so, the effects of exogenous cues could be individually tested on human neural stem/progenitor cell proliferation and neuronal network formation. Similarly, cells in 3D cultures can be transfected with plasmids for misexpression studies, a powerful tool for rapid investigation of gene function.

Our new 3D culture platform provides a novel and improved model to study the plasticity of human neural stem cells and disease states in real time. The application of customized Aβ42 peptides for optimized cellular uptake within the hydrogel-based 3D cultures indicates that our advanced culture method can recapitulate the major pathophysiology of human Aβ42 toxicity. Additionally, our gels can be used to experimentally investigate how new cells are incorporated into diseased brains, and how different sources of neural stem cells would molecularly affect the outcomes of Aβ42 toxicity (e.g.: primary versus induced neural stem cells). Individual differences between cell types and their response to disease stimuli can also be measured in a tissue-mimetic composition using our gel system. In overall, starPEG-heparin-hydrogel-based 3D human neural stem cell culture is a novel and comprehensive method for analyzing various stages of neural development and disease of the human cortex, from stem cell proliferation to neurogenesis and from neuronal maturation to integration into the circuitry, in a highly defined and controllable method in an *in vitro* environment. Our system can be expanded for examining embryonic stem cells, organoids or adult-derived cortical cells from humans, and analyze their stem cell properties or neurogenic capacity in a comparative manner to complement previous studies (Choi et al., 2014; Koutsopoulos and Zhang, 2013; Zhang et al., 2014). Beyond that, our established 3D hydrogel cultures can be expected to enable personalized medicine approaches targeting brain diseases or drug efficacy tests.

## Author contributions

C.P. and C.K. conceived and designed the experiments. M.I.C. analyzed the next generation sequencing data and performed the bioinformatics analyses. L.B., U.F., and C.W. provided the gel materials, J.F. performed AFM studies. C.P. and H.C. performed cell cultures, imaging and quantifications. P.B., H.H. and V.M helped the cell cultures. C.P. optimized the culture conditions for iPSCs and Matrigel. A.K.T. and Y.Z. provided the Amyloid peptides. X.C., S.H. and C.L.A. performed whole-cell patch clamping. C.K. wrote the manuscript, C.K., C.P., U.F., and C.W. revised the manuscript.

## Acknowledgements

This work was supported by DZNE and Helmholtz Association (VH-NG-1021, C.K.), DFG (KI1524/6, C.K.), (AN797/4-1, C.L.A.) and (CRC TR 67, CRC SFB 655, FOR/EXC999, C.W.); and BMBF (PRECIMATRIX-FKZ-03XP0083-310117, C.W.). We wish to thank M. Wagner for electrophysiology, R. Szech for image rendering, and T. Kurth for electron microscopy.

## Materials and Methods

### Cell Culture

Astrocytes isolated from the cerebral cortex at gestation week 21 were obtained from ScienCell Research Laboratory (SRL, Catalog Number 1800, Carlsbad, CA, USA) at passage one and delivered as frozen stocks. The cells (from here on primary human neural stem cells, pNSC) are certified to be negative for HIV-1, HBV, HCV, mycoplasma, bacteria, yeast, and fungi. PHCCs were seeded on conventional T75 flasks or 24-well plates and cultured in complete astrocyte medium (AM) composed of Astrocyte medium (SRL, Catalog Number 1801) supplemented with 2% fetal bovine serum (SRL, Catalog Number 0010), 1% astrocyte growth supplement (SRL, Catalog Number 1852) and 1% penicillin/streptomycin solution (SRL, Catalog Number 0503) in an incubator with a 5% CO2/95% air atmosphere at 37 °C. Human induced neural stem cells (HIP, BC1 line, Amsbio, catalog number GSC-4311, from here on iNSC) were seeded in Geltrex pre-coated cultureware in complete AM as described in the above paragraph.

### Generation of starPEG-Heparin hydrogels and cell encapsulation

StarPEG-heparin hydrogels were generated as previously described (Maitz et al., 2013; Wieduwild et al., 2013) with the following modifications: pNSCs or iNSCs were collected from culture flasks using Accutase (Invitrogen, CA, USA). After centrifugation (271 g for 10 minutes), cells were resuspended in PBS at a concentration of 8 × 10^6^ cells per ml. For each hydrogel, we first resuspended the cells in 5 μl of PBS, then added 5 μl Heparin maleimide conjugate solution (90 μg/μl in PBS) and 10 μl starPEG-MMP-peptide conjugate solution (Tsurkan et al., 2011) to a final volume of 20 μl and a cell density of 2 × 10^6^ cells/ml. Next, the 20 μl droplet was placed on a Parafilm sheet for approximately two minutes until it began to gelate. The gels were placed in 24-well plates, and each well contained 1 ml of astrocyte culture medium (AM) (Supplementary Movie 8). The gels were cultured and incubated in 5% CO2/95% air at 37 °C until the desired time points (1 week, 2 weeks, and 3 weeks).

### Generation of Matrigel cultures

For the generation of Matrigel cultures, we used BD Matrigel (catalog number: 356234). Prior to any cell culture work and use of the Matrigel, pipette tips and Eppendorf tubes has been frozen at −20 °C according to manufacturer’s instruction following the “thick gel method”. The BD Matrigel has been thawed overnight on ice at 4 °C. pNSCs were collected from culture flasks using Accutase (Invitrogen). After centrifugation (271 g for 10 minutes), pNSCs were re-suspended in BD Matrigel in concentration 2×10^6^ cells per ml. Droplets of the cell/Matrigel mix were placed in the bottom of culture and then waiting to solidify at 37 °C. Then, cell medium (SRL, Catalog Number 1801) added and the gels cultured for 3 weeks (Supplementary Movie 8). Cell medium changed the day after the generation of gels and then every other day.

### Synthesis of Amyloid peptides

The peptides were synthesized as previously described (Bhattarai et al., 2017a; Bhattarai et al., 2016; Bhattarai et al., 2017b; Kizil et al., 2015; Wieduwild et al., 2013). For peptide synthesis, all of the required chemicals were purchased from IRIS Biotech GmbH (Marktredwitz, Germany). Acetonitrile (for UPLC/LCMS), dichloromethane (DCM), diethylether, dimethyl sulfoxide (DMSO), formic acid (FA), trifluoroacetic acid (TFA), and triisopropylsilane (TIS) were purchased from MERCK KGaA (Darmstadt, Germany). Acetic anhydride and N-methylmorpholine (NMM) were purchased from Sigma-Aldrich Co. LLC (St. Louis, MO, USA). Dithiothreitol (DTT) was obtained from Prolab VWR International, LCC (Radnor, PA, USA). Acetonitrile (for HPLC) was purchased from TH Geyer (Renningen, Germany). 5(6)-Carboxyfluorescein was purchased from Acros Organics (Fisher Scientific Company LLC). The TentaGel S RAM Fmoc rink amide resin was purchased from RappPolymere GmbH (Tübingen, Germany). The peptide synthesis columns and syringes with included filters were purchased from Intavis AG (Cologne, Germany). Water was obtained from a Milli-Q water purifier (Milli-Q Advantage A10, EMD Millipore Corporation, Billerica, MA, USA) with a LCPAK0001 Milli-Q filter. The polytetrafluoroethylene (PTFE) filter, polyvinylidene fluoride (PVDF) syringe filter, and filter holder were purchased from Sartorius Stedtim (Aubagne, France). Aβ42 peptides were prepared using standard 9-fluorenylmethoxycarbonyl (Fmoc) chemistry with 2-(1H-benzotriazol-1-yl)-1,1,3,3-tetramethyluronoium hexafluorophosphate (HBTU) activation on an automated solid-phase peptide synthesizer (ResPep SL, Intavis)(Wieduwild et al., 2013; Zhang et al., 2002). Each amino acid was coupled twice at 5-fold excess followed by capping the non-reacted amino groups with acetic anhydride to achieve high quality synthesis. Upon completion of the peptide synthesis, 5(6)-carboxyfluorescein was coupled to the N-terminus using HBTU as the coupling reagent. The peptide was then cleaved from the resin with TFA/TIS/water/DTT (90(v/v):5(v/v):2.5(v/v):2.5(m/v)) for 2 hours. The product was precipitated and washed with ice-cold diethyl ether.

The peptide was dissolved in Milli-Q water, and peptide purification was performed via reverse-phase high-pressure liquid chromatography (HPLC) on a semi-preparative HPLC (Waters) equipped with a semi-preparative column (PolymerX RP-1, 250 × 10 mm, Phenomenex). The peptide was eluted from the column by applying a gradient of 5% to 100% solvent B over 30 min at 20 ml/min, in which solvent A is 0.1% TFA in water and solvent B is 0.1% TFA and 5% water in acetonitrile.

Purity was confirmed on an analytical reverse phase ultra-high pressure liquid chromatograph (UPLC Aquity with UV Detector) equipped with an analytical C18 column (Acquity UPLC BEH C18, bead size 1.7 μm, 50 × 2.1 mm) using an isocratic gradient and electrospray ionization mass spectrometry (ESI-MS) (Acquity TQ Detector).

### Aβ42 treatment

The Aβ42 treatment was performed 24 h post-thaw for a period of 48 h at 10 μM final concentration for pNSCs and iNSCs in 3D cultures. Amyloid plaques remain in the culture attached to the cells throughout the cultures. The medium was removed and the cells were washed with PBS after 48 h Aβ42 treatment. Then, the cells were collected using Accutase, counted and centrifuged at 271 g for 10 min. The cell pellet was resuspended in PBS to obtain at 8 × 10^6^ cells/ml. This cell suspension was mixed with an equal volume of 6.00 mM heparin maleimide conjugate in PBS to obtain a 3.00 mM heparin maleimide conjugate-cell suspension mix at 4.0 × 10^6^ cells/ml. 10 μl of this mix was combined with 10 μl of 2.25 mM starPEG-MMP-peptide conjugate solution in PBS (Tsurkan et al., 2011), quickly triturated a few times, and the resulting 20 μl volume was pipetted onto a Parafilm sheet forming a droplet. This droplet was allowed to gelate for about 2 min resulting in formation of a 20 μl hydrogel containing 40,000 cells (2.0 × 10^6^ cells/ml) at a final concentration of 1.50 mM heparin maleimide conjugate and 1.12 mM starPEG-MMP-peptide conjugate. The gels were then placed in 0.75 ml of AM culture medium per well in 24-well plates, and incubated in 5% CO2/95% air at 37 °C until the desired time points (1 week, 2 weeks, and 3 weeks). The medium was changed 3 times a week throughout the incubation period. 2D pNSC cultures were treated with 2 μM Aβ42 in the culture medium for 48 hours between second and fourth days of culture. The medium is removed and cells are washed with cell culture medium twice. Amyloid plaques remain in the culture attached to the cells throughout the cultures. New medium is added and cultures were continued.

### Transplantation

Primary human neural stem cells (pNSCs) were cultured for two days and treated with TOPRO-3 (Thermo Scientific) for 1 hour before transplantation. After a washing step, the cells were harvested with Accutase and re-suspended in astrocyte cell culture medium at 3 × 10^5^ cells per ml. With a pipette tip, 2 μl of the cell suspension was injected into the center of the hydrogel. After a week of culture, the hydrogels were fixed and processed for immunocytochemistry.

### Immunocytochemistry

All of the hydrogels were fixed with ice-cold 4% paraformaldehyde and incubated for 1.5 hours at room temperature followed by washing in PBS overnight at 4 °C. For immunocytochemistry, the hydrogels were blocked and permeabilized in blocking solution for 4 hours at room temperature. For BrdU-treatment, the gels were incubated with 2 M HCl for 20 minutes at 37 °C followed by three washes in PBS (2 hours each). EdU staining was performed according to the manufacturer’s protocol (Life Technologies, C10638) using a 1-hour incubation step. The hydrogels were incubated with primary antibodies (Supplementary Table 1) in blocking solution overnight at 4 °C. The gels were washed for two subsequent days at 4 °C, with occasional changes of the PBS. After washing, the gels were incubated with the secondary antibodies (1:500 in blocking solution) at room temperature for 6 hours. After 3 washing steps of 2 hours each, DAPI staining was performed (1:3000 in PBS, 2 hours at room temperature). Immunostaining for SOX2 (Santa Cruz Biotechnology, 1:100), SATB2 (Abcam, 1:300), ASCL1 (Neuromics, 1:100), CTIP2 (Abcam, 1:100), EEA1 (Abcam, 1:500), neurofilament (NF-M+H+L) (Life Technologies, 1:500), TUBB3 (R&D Systems, 1:500), CASP3 (Santa Cruz Biotechnology, 1:500), MKI67 (Abcam, 1:1000), acetylated tubulin (Sigma, 1:500), SYN (Millipore, 1:500), GFAP (Novex, 1:500), DCX (Novex, 1:300), MAPT (Abcam, 1:500), Aβ42 (Cell Signaling Technology, host: Rabbit, 1:500), Reelin (Abcam, 1:500), FOXO1 (Thermo Scientific, 1:500), FOXP2 (R&D, 1:200), CRYM (Thermo Fischer, 1:50), PSD95 (Thermo Fischer, 1:300), DBX1 (Abcam, 1:300), VGLUT1 (Thermo Fischer, 1:500), BrdU (AdB Serotec, 1:500) was performed. All of the secondary antibodies were conjugated to AlexaFluor dyes (Life Technologies).

### Fluorescent imaging

For the hydrogels, fluorescent imaging was performed using a Leica SP5 inverted Laser Scanning Confocal microscope. The hydrogels were placed in glass bottom Petri dishes. Sixty microliters of PBS were added on top of the hydrogels to avoid desiccation. The Z-stacks were captured using a 25x water immersion lens. Every Z-stack had a z-distance of 500 μm. Monolayers were imaged using an inverted Zeiss Apotome 2 microscope.

### Histological analyses

For Gallyas silver staining, the 3D hydrogels were cryo-frozen and sequentially incubated in 5% periodic acid (5 minutes), an alkaline silver iodide solution (1 minute), acetic acid (3 minutes), 0.1% gold chloride (5 minutes), 1% sodium thiosulfate (5 minutes), and 2.5% aluminum sulfate (1 minutes), with intermittent washes with distilled water. For Thioflavin S staining, samples were incubated in 1% Thioflavin S (8 minutes), absolute ethanol (3 minutes), and DAPI (10 minutes).

### Transfection with GCaMP6f plasmids and calcium imaging

TurboFectin 8.0 reagent (OriGene, Cat# TF81001) was used to transfect adherent (2D cultures) and encapsulated (3D cultures) cells with 700 μg plasmid per reaction in 1 ml of cell growth medium. The pGP-CMV-GCaMP6f plasmid was a gift from Douglas Kim (Addgene plasmid # 40755)(Chen et al., 2013). The images were captured using a Leica SP5 inverted Laser Scanning Confocal microscope in resonant scanner mode with photon counting. Images were acquired every 100 milliseconds. Analysis of the calcium image spectrum was performed with the Leica LAS AF software by using region of interests (ROIs) and photon counting.

### Patch clamp recordings

Single neurons were recorded in Artificial Cerebrospinal Fluid (ACSF) (119 mM NaCl, 2.5 mM KCl, 2 mM CaCl2, 1.3 mM MgCl2, 1 mM NaH2PO4, and 10 mM glucose, pH 7.3) and patched with nerve solution (125 mM K+-gluconate, 0.1 mM CaCl2, 0.6 mM MgCl2, 8 mM NaCl, 1 mM EGTA, 0.01 mM HEPES, and 4 mM Na-ATP, pH 7.23) The Whole-cell patch recordings were assessed using a HEKA set up and Pulse program. Membrane voltage resistance was held at −80 mV with the pipette resistance of 4-6 MOhms. For measurements of K+ and Na+ currents, test pulses were applied in 80 ms durations from −80 mV to 30 mV every 2 s. All experiments were done at 20-23 °C.

### Electron microscopy

For electron microscopy, the hydrogel-embedded cells were fixed in modified Karnovsky’s fixative (2% glutaraldehyde + 2% paraformaldehyde in 50 mM HEPES) at least overnight at 4 °C. The samples were washed 2x in 100 mM HEPES and 2x in water and postfixed in a 2% aqueous OsO4 solution containing 1.5% potassium ferrocyanide and 2 mM CaCl_2_ for 30 min on ice. Next, washes in water, 1% thiocarbohydrazide in water (20 minutes at room temperature), water, and a second osmium contrasting step in 2% OsO_4_/water (30 minutes on ice). After several washes in water, the samples were *en bloc* contrasted with 1% uranyl acetate/water for 2 hours on ice, washed again in water, dehydrated in a graded series of ethanol/water up to 100% ethanol, and infiltrated with Epon 812 (Epon/ethanol mixtures: 1:3, 1:1, 3:1 for 1.5 hours each, pure Epon overnight, and pure Epon for 5 hours). The samples were embedded in flat embedding molds and cured overnight at 65 °C. Ultrathin sections were prepared with a Leica UC6 ultramicrotome (Leica Microsystems, Vienna, Austria), collected on Formvar-coated slot grids and stained with lead citrate and uranyl acetate as previously described(Venable and Coggeshall, 1965).

For CLEM, cells that were embedded in the hydrogels with the fluorescein-labeled peptide were fixed with 4% paraformaldehyde in 100 mM phosphate buffer (PB). After several washes in water, the samples were dehydrated in 50% (15 minutes at 4 °C), 70%, 90%, and 100% acetone (45 minutes each at −25 °C) and incubated with LR Gold (London Resin Company, Reading, UK) solutions of 33% and 66% LR Gold/acetone, pure LR Gold (1 hour each at −25 °C), and LR Gold + 0.1% benzil (1 hour, overnight at −25 °C). Finally, the samples were transferred to LR Gold-containing 1% benzil and polymerized using the UV lamp of the Leica AFS2 freeze substitution unit (Leica Microsystems, Vienna, Austria) for 48 hours at −25 °C. Ultrathin sections were mounted on Formvar-coated EM grids, stained with DAPI, imaged with a wide field fluorescence microscope, washed, and contrasted with 1% uranyl acetate for EM as previously described(Fabig et al., 2012). Contrasted ultrathin sections were analyzed on a FEI Morgagni D268 (FEI, Eindhoven, The Netherlands) or a Jeol JEM1400 Plus at 80 kV acceleration voltage.

### Atomic force microscopy

Atomic force microscopy (AFM) was performed to determine the mechanical properties of the gels. Briefly, AFM measurements were collected at 37 °C using a Nanowizard II AFM (JPK Instruments, Berlin, Germany). Tipless silicon nitride cantilevers with a nominal spring constant of 80 mN*m-1 (PNP-TR-TL-Au; Nanoworld) were used. The cantilevers were modified with silica beads (Æ10 μm, Kisker Biotec GmbH), as previously described (Bray et al., 2015). Force-distance curves were acquired in closed loop, constant height mode using a 3 nN contact force and a 5 μm/s approach/retract velocity. Each data set was generated by probing a minimum of 70 different spots on each sample. The data processing software provided by the AFM manufacturer (JPK Instruments) was used to extract the Young’s Modulus E from the approach force-distance curves.

### RNA Isolation

RNA isolation from 2D cell culture was performed by TriZol (Invitrogen). Total RNA isolation from 3D gels was performed by Norgen Total RNA isolation Kit (Cat#17200). 5 gels were lysed in 1ml RL buffer with 10 μL β-mercaptoethanol, and after centrifugation at 12,000g for 7 min at room temperature, the supernatant was collected in a new Eppendorf and mixed with absolute Ethanol. The remaining steps performed as previously described (Bhattarai et al., 2016).

### Next generation sequencing of whole transcriptome

cDNA libraries were prepared by following the protocol for NEBNext^®^ Ultra I Directional RNA Library Prep Kit. This involves the following steps: mRNA isolation via poly(A)+ selection and fragmentation, first strand and second strand cDNA synthesis, purification using the Agencourt^®^ AMPure^®^ Kit and end repair/dA-tailing of cDNA. Adapters were ligated to the dA-tailed cDNA, followed by an size selection using AMPure XP Beads. Indexing of the library constructs was done with illumina^®^ index primer during the following PCR amplification using NEBNext^®^ Q5 2X PCR Master Mix. Lastly, libraries were purified using the Agencourt^®^ AMPure^®^ Kit. Libraries were pooled and sequenced on an illumina^®^ NextSeq 500 system, resulting in ca. 27 – 38 million 75 bp single-end reads. All protocols are performed according to the manufacturers’ instructions.

### Data Analysis

The reads in fastq files were aligned to the human genome (hg19/GRCh38) with gsnap (version 2016-09-23) (Wu et al., 2016), and featureCounts (v1.5.3) (Liao et al., 2013, 2014) was used to assign reads to each gene using Ensembl version 90 [Homo_sapiens.GRCh38.90.gtf]. DESeq2 (1.18.0) (Love et al., 2014) was used to normalize the reads, calculate fold changes and p-values. 2-fold change and padj value of less than 0.1 were used to identify differentially expressed genes. For KEGG pathway analysis GOstats (2.44.0) (Falcon and Gentleman, 2007), GOSeq (1.30.0) (Young et al., 2010), and clusterProfiler (3.6.0) (Yu et al., 2012) were used. topGO (2.30.0) (Alexa et al., 2006) was used for GO analysis and pathview (1.18.0) (Luo and Brouwer, 2013) was used for drawing KEGG pathway. All data analysis pipeline scripts were written in R in our lab, and are available upon request.

### Sequencing datasets

All deep sequencing experiments (pNSC 2D, pNSC 3D, iNSC 3D control and Amyloid-beta42-treated) can be found under the GEO accession number GSE78117.

### Image analysis and statistics

The 3D reconstructions of hydrogel images and videos were generated using Arivis 4D software. Images from monolayers were processed using Zeiss ZEN software. The statistical analyses were performed using GraphPad Prism and two-tailed Student’s t-tests. The levels of significance were *: p ≤ 0.05, **: p ≤ 0.01, and ***: p ≤ 0.001. In all graphs, means ± standard deviations are shown.

The effect size was calculated using G-Power, and the sample size was estimated with n-Query. The data conforms to normal distribution as determined by Pearson’s chi-squared test. The variations between the samples are similar as determined by variance estimation using Microsoft Excel software. For 3D gels, 9 gels were used for quantifications (3 technical replicates in every experiment, and 3 experiments as biological replicates). All experiments were replicated many times in the laboratory and results were confirmed independently (80–120 gels were qualitatively analyzed to check the consistency of the results for every individual experiment).

### Generation of skeletonized networks and quantification

To examine the axons of neural cells, the length and branching were obtained by thinning binary images to a skeleton, which was performed in all three dimensions. In detail, the raw images were processed with a Gaussian filter and then with the tubeness filter to enhance linear structures. Then, an automatic threshold was applied, followed by several morphological operations to facilitate the skeletonization. Fiji software (www.fiji.sc) was used for image processing. Skeletons were quantified using KNIME freeware (Supplementary Figure 4).

## Supplementary Information

Supplementary Figure 1-5 Supplementary Movies 1-8 Supplementary Dataset 1-12

**Supplementary Figure 1.**
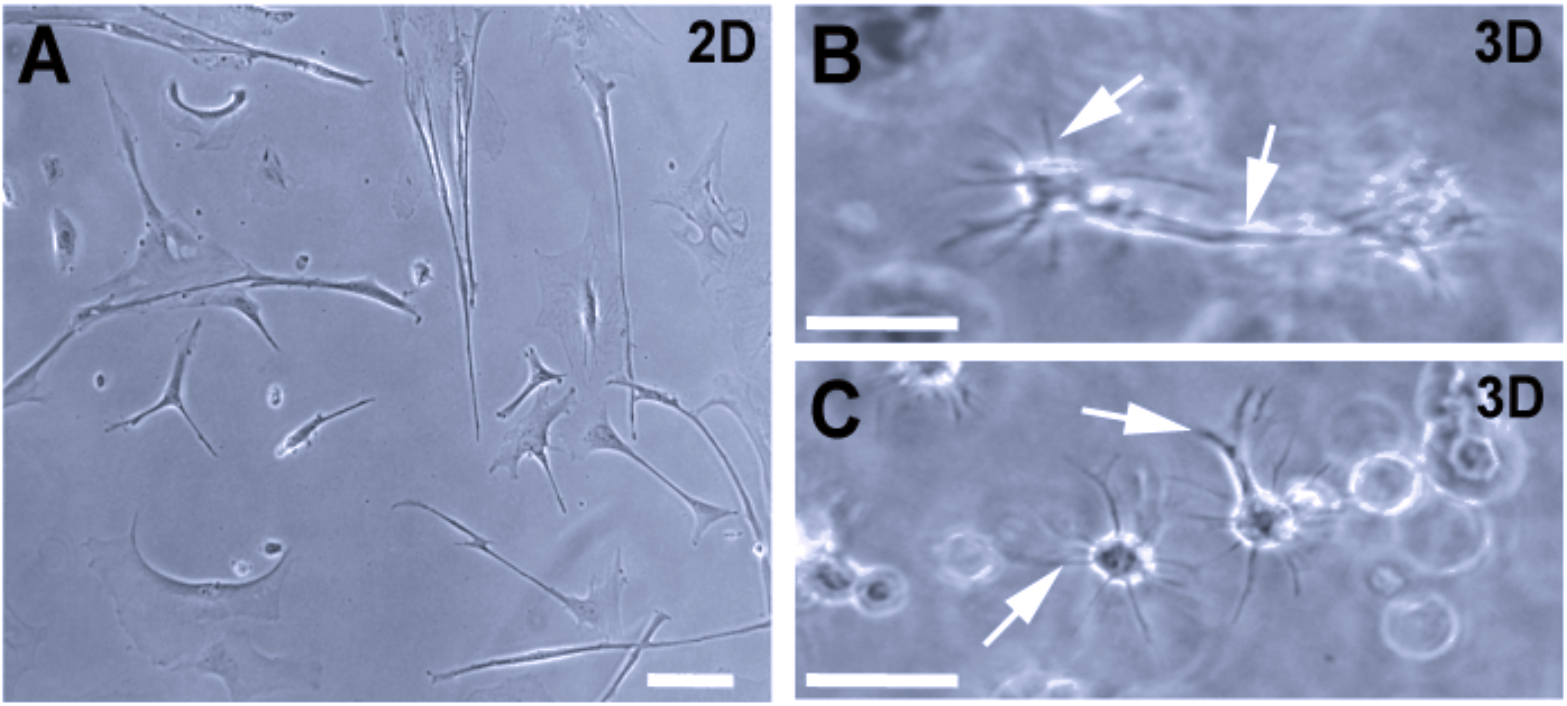
Formation of 3D topology and arborizations in starPEG-Heparin gels. (A) Culture of primary human NSCs in 2D. (B,C) Culture of primary human NSCs in 3D. Note the arborized morphology and cellular processes reminiscent of in vivo (white arrows). Scale bars 25 μm. Related to Figure 1.

**Supplementary Figure 2.**
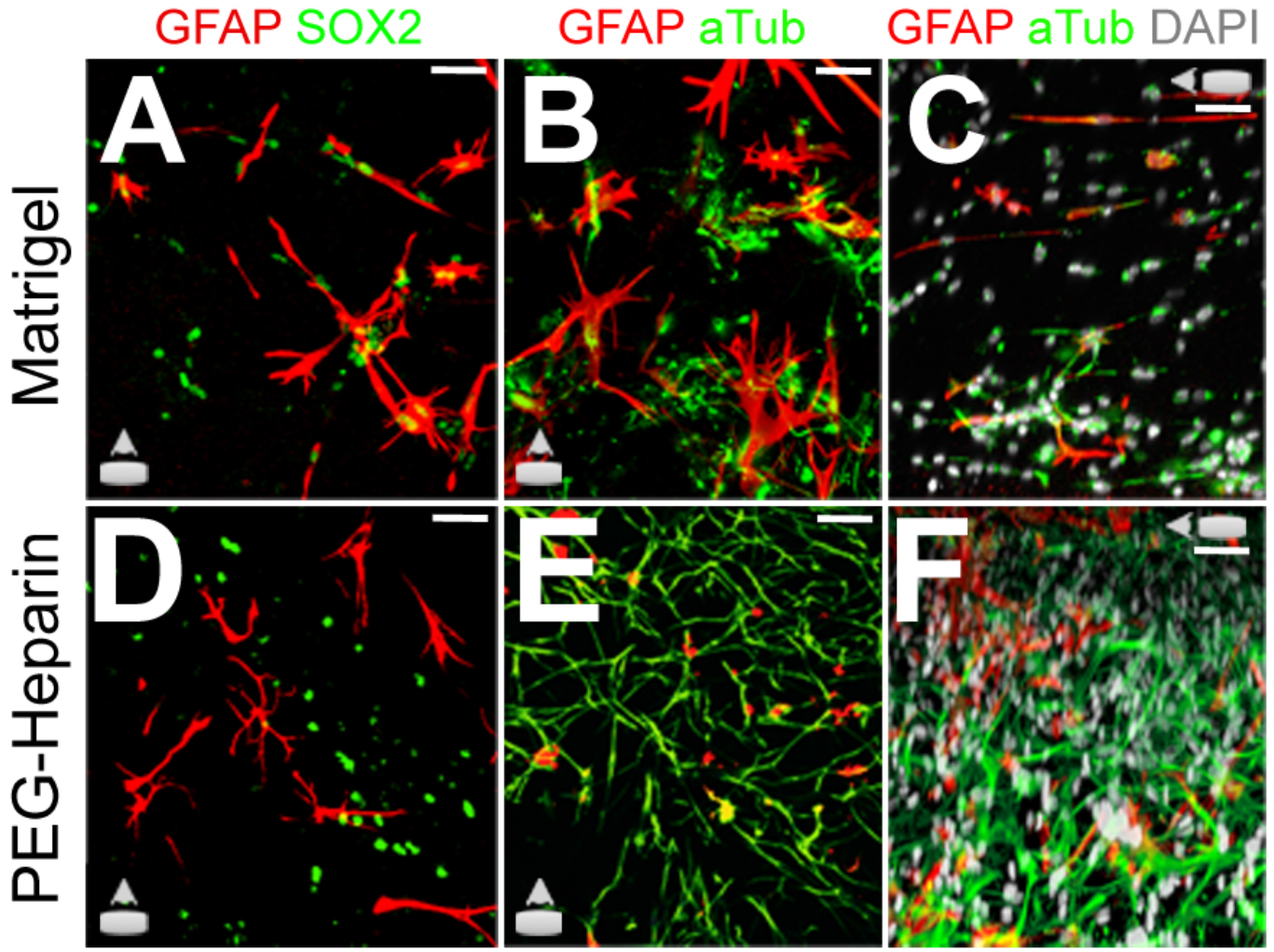
Comparison of Matrigel and starPEG-Heparin cultures. (A) Immunostaining for GFAP and SOX2 in Matrigel cultures. (B) Immunostaining GFAP and Acetylated tubulin in Matrigel cultures Z-view. (C) Immunostaining GFAP and Acetylated tubulin in Matrigel cultures X-view. (D) Immunostaining for GFAP and SOX2 in starPEG-Heparin cultures. (E) Immunostaining GFAP and Acetylated tubulin in starPEG-Heparin cultures Z-view. (F) Immunostaining GFAP and Acetylated tubulin in starPEG-Heparin cultures X-view. Scale bars: 20 μm. All gels are 3 weeks of culture.

**Supplementary Figure 3.**
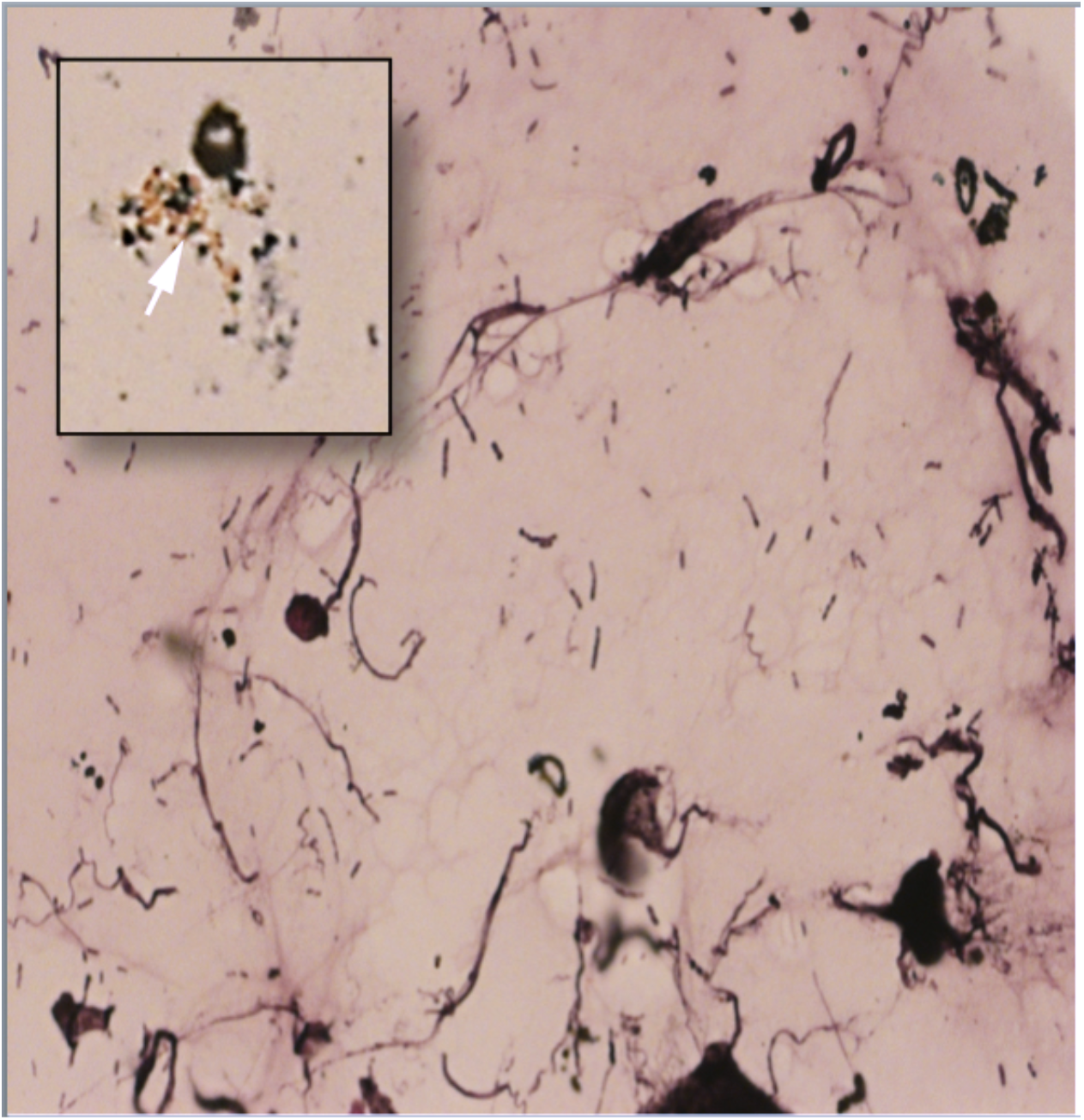
*Formation of neurofibrillary tangles and senile plaques in 3D cultures of primary human NSCs*. Image shows Gallyas silver impregnation staining for neurofibrillary tangles in 3D cultures. Inset shows the senile plaques (white arrow).

**Supplementary Figure 4.**
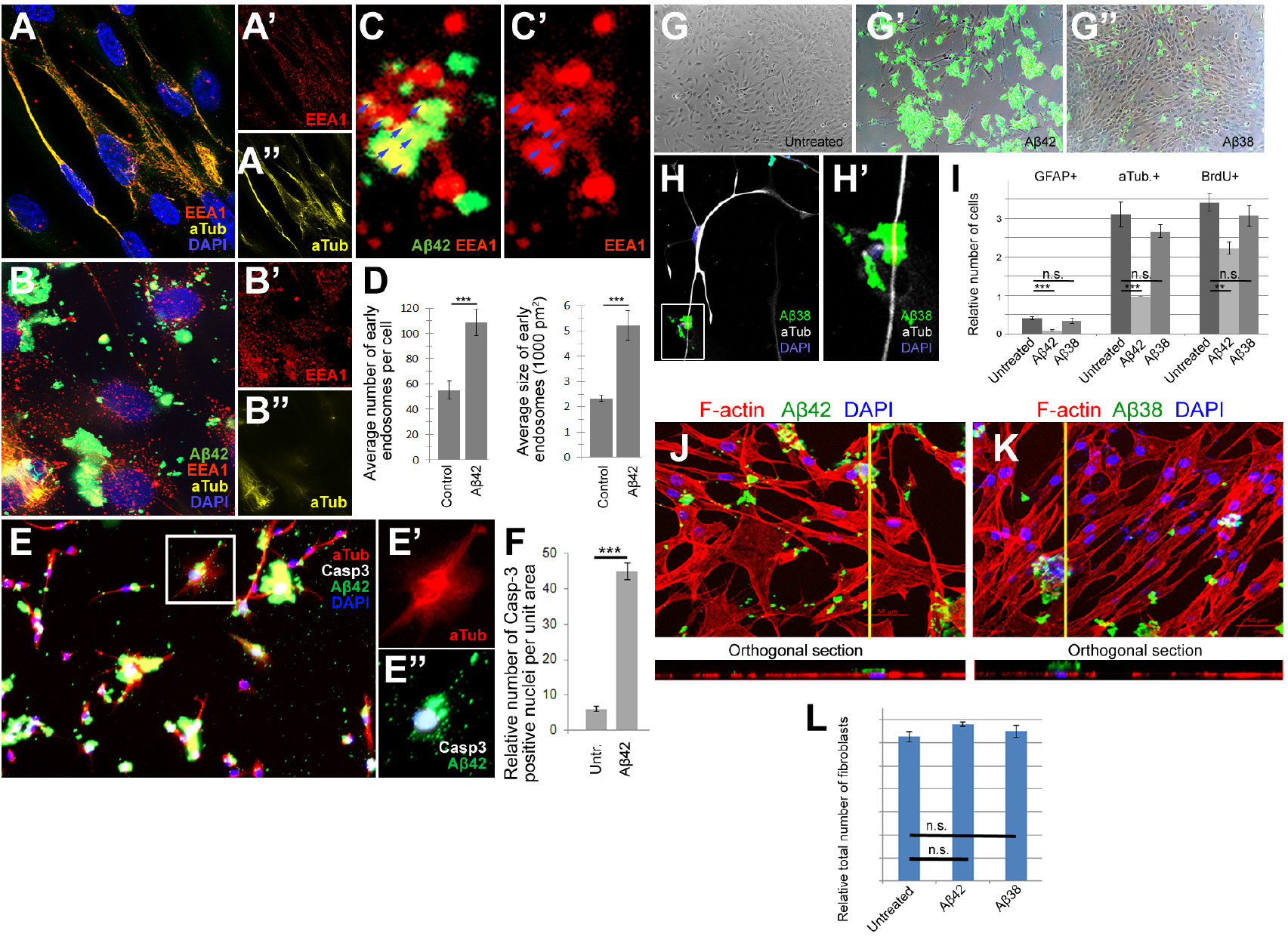
Amyloid aggregation dynamics. (A) Immunostaining for acetylated tubulin and early endosomes (EEA1) in control cultures. (A’, A’’) Individual fluorescence channels for EEA1 and acetylated tubulin in control cultures. (B) Immunostaining for acetylated tubulin and early endosomes (EEA1) in Aβ42-treated cultures. (B’, B’’) Individual fluorescence channels for EEA1 and acetylated tubulin in Aβ42-treated PHCCs. (C, C’) Co-localization of EEA1 and Aβ42. (D) Quantification of the average number of early endosomes per cell and the average size of early endosomes per cell in control and Aβ42-treated cultures. (E) Immunostaining of Aβ42-treated cells for acetylated tubulin (red), Aβ42 (green) and caspase-3 (white). (E’) Individual fluorescence channel for acetylated tubulin. (E’’) Fluorescence channels for Aβ42 and Caspase-3. (F) Quantification of Caspase-3-positive cells in control and Aβ42-treated cultures. (G) Bright field image of control cultures. (G’) Bright field image of Aβ42-treated cultures. (G’’) Bright field image of Aβ38-treated cultures. (H) Confocal image of acetylated tubulin immunostaining on Aβ38-treated cultures. (H’) Close-up of a region from H showing cells treated with Aβ38. (I) Quantification of the relative number of untreated, Aβ42-treated and Aβ38-treated cells immunoreactive for GFAP, acetylated tubulin or BrdU. (J) Confocal image over the z-axis and orthogonal section over the y-axis of Aβ42-treated human-derived fibroblasts. F-actin was stained with phalloidin, and DNA was stained with DAPI. (K) Confocal image over the z-axis and orthogonal section over the y-axis of Aβ38-treated human-derived fibroblasts. F-actin was stained with phalloidin, and DNA was stained with DAPI. (L) Quantification of the relative total number of fibroblasts in control, Aβ42-treated and Aβ38-treated samples. Scale bars 20 μm. Related to Figure 4.

**Supplementary Figure 5.**
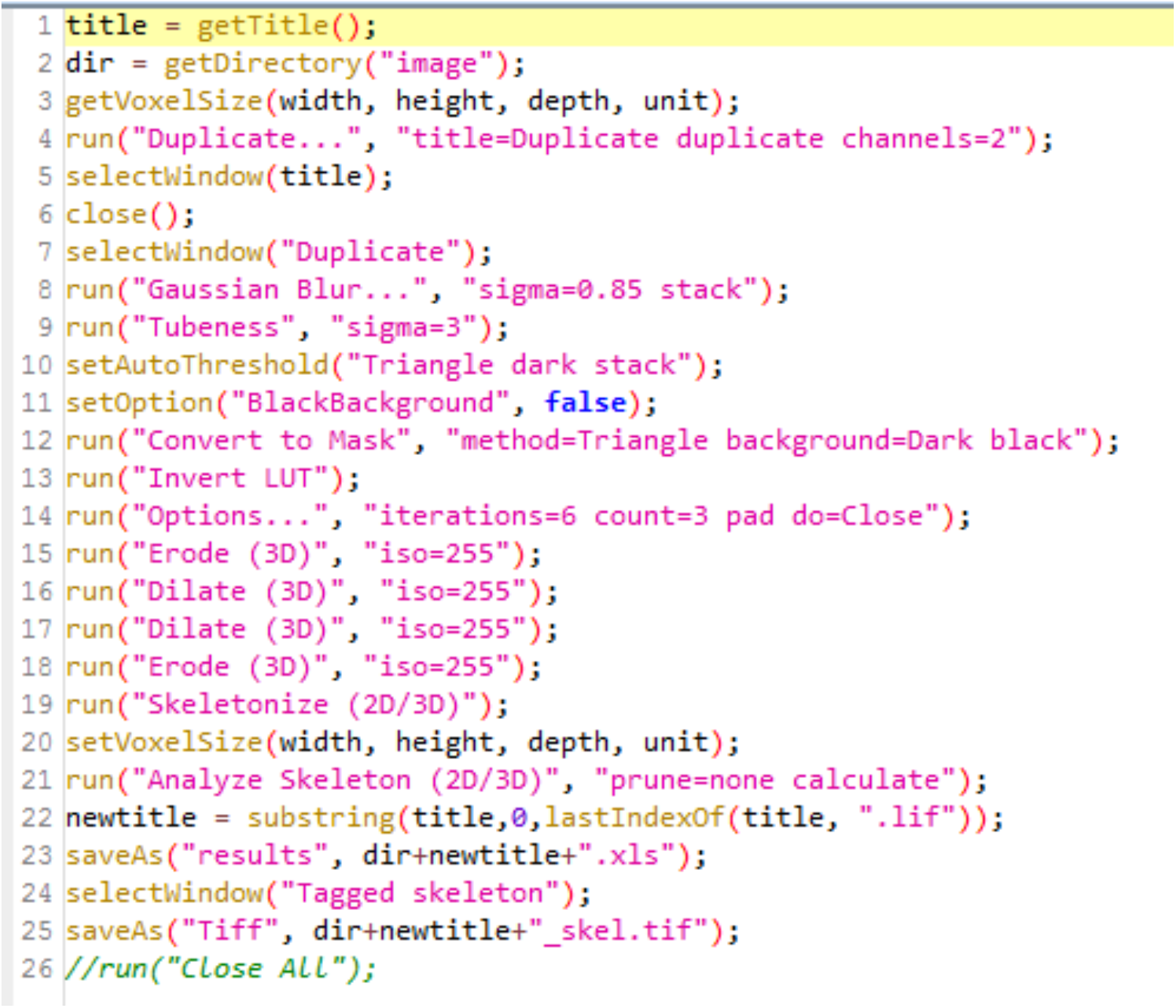
*Script for KNIME software*

## Supplementary Movies

### Supplementary Movie 1

#### Neuronal network in pNSC cultures

3D representation of a 0.18 mm^3^ portion of a control hydrogel containing networks of neurons as assessed by acetylated tubulin staining (cyan). Related to Figure 1.

### Supplementary Movie 2

#### Neuronal network in iNSC cultures

3D reconstruction of iPSC-derived NSC cultures in PEG-Heparin gels. Cyan: acetylated tubulin, white: DAPI, red: GFAP. Related to Figure 1.

### Supplementary Movie 3

#### Calcium activity

Encapsulated primary human cortical astrocytes in PEG-HEP gels transfected with the GCaMP6f Calcium sensor exhibit green fluorescence in response to calcium influx after treatment with glutamate. Related to Figure 1.

### Supplementary Movie 4

#### Dystrophic axons

A TUBB3-positive neuron (cyan) with retracted processes, dystrophic axonal ends, and Aβ42 aggregation (green) is shown in a 3D reconstruction. DAPI is shown in white. Related to Figure 2.

### Supplementary Movie 5

#### 3D reconstruction of a neuron with hyperphosphorylated Tau

A 3D reconstruction of primary human cortical astrocyte cultures after 3 weeks in Aβ42-treated gels and staining for Aβ42 (green), DAPI (white) and phosphorylated Tau (AT180, red). Note that neurons exhibit staining for hyperphosphorylated Tau. Related to Figure 2.

### Supplementary Movie 6

#### Transplantation in control gels

3D reconstruction of a labeled neuron (red nuclei, transplanted) that established connections to the existing neurons in the control gels. Cyan: acetylated tubulin, white: DAPI, red: TOPRO3. Related to Figure 2.

### Supplementary Movie 7

#### Transplantation in Amyloid-containing gel

3D reconstruction of a labeled neuron (red nuclei, transplanted) that was unable to make connections with the existing, already dystrophic, neurons in amyloid-containing gels. Green: Aβ42, cyan: acetylated tubulin, white: DAPI, red: TOPRO3. Related to Figure 2.

### Supplementary Movie 8

#### Comparison of gel preparation

Comparison of preparation of starPEG-Heparin and Matrigel cultures of primary human astrocytes. Related to Figure 2.

## Supplementary Datasets

### Supplementary Dataset 1

Differential expression analyses between pNSC 3D cultures and pNSC 2D cultures.

### Supplementary Dataset 2

Elim Fischer cellular component analyses of differentially expressed genes between pNSC 3D cultures and pNSC 2D cultures.

### Supplementary Dataset 3

KEGG pathway analysis of differentially expressed genes between pNSC 3D cultures and pNSC 2D cultures.

### Supplementary Dataset 4

Differential expression analyses between iNSC 3D cultures and pNSC 3D cultures.

### Supplementary Dataset 5

Elim Fischer cellular component analyses of differentially expressed genes between iNSC 3D cultures and pNSC 3D cultures.

### Supplementary Dataset 6

KEGG pathway analysis of differentially expressed genes between iNSC 3D cultures and pNSC 3D cultures.

### Supplementary Dataset 7

Differential expression analyses upon Aβ42 in 3D iNSC cultures.

### Supplementary Dataset 8

Differential expression analyses upon Aβ42 in 3D pNSC cultures.

### Supplementary Dataset 9

Elim Fischer cellular component analyses of differentially expressed genes in Aβ42-treated 3D iNSC cultures.

### Supplementary Dataset 10

KEGG pathway analysis of differentially expressed genes in Aβ42-treated 3D iNSC cultures.

### Supplementary Dataset 11

Elim Fischer cellular component analyses of differentially expressed genes in Aβ42-treated 3D pNSC cultures.

### Supplementary Dataset 12

KEGG pathway analysis of differentially expressed genes in Aβ42-treated 3D pNSC cultures.

## References

Agrawal, S.M., Lau, L., and Yong, V.W. (2008). MMPs in the central nervous system: where the good guys go bad. Semin Cell Dev Biol 19, 42–51.

Alexa, A., Rahnenfuhrer, J., and Lengauer, T. (2006). Improved scoring of functional groups from gene expression data by decorrelating GO graph structure. Bioinformatics 22, 1600–1607.

Alvarez-Buylla, A., Seri, B., and Doetsch, F. (2002). Identification of neural stem cells in the adult vertebrate brain. Brain Res Bull 57, 751–758.

Bhattarai, P., Thomas, A.K., Cosacak, M.I., Papadimitriou, C., Mashkaryan, V., Zhang, Y., and Kizil, C. (2017a). Modeling Amyloid-β42 Toxicity and Neurodegeneration in Adult Zebrafish Brain. Journal of Visualized Experiments 128.

Bhattarai, P., Thomas, A.K., Papadimitriou, C., Cosacak, M.I., Mashkaryan, V., Froc, C., Kurth, T., Dahl, A., Zhang, Y., and Kizil, C. (2016). IL4/STAT6 signaling activates neural stem cell proliferation and neurogenesis upon Amyloid-β42 aggregation in adult zebrafish brain. Cell Reports 17, 941–948.

Bhattarai, P., Thomas, A.K., Zhang, Y., and Kizil, C. (2017b). The effects of aging on Amyloid-β42-induced neurodegeneration and regeneration in adult zebrafish brain. Neurogenesis.

Bray, L.J., Binner, M., Holzheu, A., Friedrichs, J., Freudenberg, U., Hutmacher, D. W., and Werner, C. (2015). Multi-parametric hydrogels support 3D in vitro bioengineered microenvironment models of tumour angiogenesis. Biomaterials 53, 609–620.

Capila, I., and Linhardt, R.J. (2002). Heparin-protein interactions. Angew Chem Int Ed Engl 41, 391–412.

Capilla, I., and Linhardt, R.J. (2002). Heparin-protein interactions,. Angew Chem Int Ed Engl 41, 391–412.

Chen, T.W., Wardill, T.J., Sun, Y., Pulver, S.R., Renninger, S.L., Baohan, A., Schreiter, E.R., Kerr, R.A., Orger, M.B., Jayaraman, V., et al. (2013). Ultrasensitive fluorescent proteins for imaging neuronal activity. Nature 499, 295–300.

Choi, S.H., Kim, Y.H., Hebisch, M., Sliwinski, C., Lee, S., D’Avanzo, C., Chen, H., Hooli, B., Asselin, C., Muffat, J., et al. (2014). A three-dimensional human neural cell culture model of Alzheimer’s disease. Nature 515, 274–278.

Chwalek, K., Bray, L.J., and Werner, C. (2014). Tissue-engineered 3D tumor angiogenesis models: Potential technologies for anti-cancer drug discovery. Adv Drug Deliv Rev.

Chwalek, K., Levental, K.R., Tsurkan, M.V., Zieris, A., Freudenberg, U., and Werner, C. (2011). Two-tier hydrogel degradation to boost endothelial cell morphogenesis. Biomaterials 32, 9649–9657.

Cosacak, M.I., Papadimitriou, C., and Kizil, C. (2015). Regeneration, Plasticity, and Induced Molecular Programs in Adult Zebrafish Brain. Biomed Res Int 2015:769763.

Demars, M., Hu, Y.S., Gadadhar, A., and Lazarov, O. (2010). Impaired neurogenesis is an early event in the etiology of familial Alzheimer’s disease in transgenic mice. J Neurosci Res 88, 2103–2117.

Discher, D.E., Mooney, D.J., and Zandstra, P.W. (2009). Growth factors, matrices, and forces combine and control stem cells. Science 324, 1673–1677.

Ermini, F.V., Grathwohl, S., Radde, R., Yamaguchi, M., Staufenbiel, M., Palmer, T.D., and Jucker, M. (2008). Neurogenesis and alterations of neural stem cells in mouse models of cerebral amyloidosis. Am J Pathol 172, 1520–1528.

Esler, W.P., and Wolfe, M.S. (2001). A portrait of Alzheimer secretases--new features and familiar faces. Science 293, 1449–1454.

Fabig, G., Kretschmar, S., Weiche, S., Eberle, D., Ader, M., and Kurth, T. (2012). Labeling of ultrathin resin sections for correlative light and electron microscopy. Methods Cell Biol 111, 75–93.

Falcon, S., and Gentleman, R. (2007). Using GOstats to test gene lists for GO term association. Bioinformatics 23, 257–258.

Fatehullah, A., Tan, S.H., and Barker, N. (2016). Organoids as an in vitro model of human development and disease. Nat Cell Biol 18, 246–254.

Freudenberg, U., Sommer, J.U., Levental, K.R., Welzel, P.B., Zieris, A., Chwalek, K., Schneider, K., Prokoph, S., Prewitz, M., Dockhorn, R., et al. (2012). Using Mean Field Theory to Guide Biofunctional Materials Design. Adv Funct Mater 22, 1391–1398.

Gage, F.H. (2000). Mammalian neural stem cells. Science 287, 1433–1438.

Gage, F.H., and Temple, S. (2013). Neural stem cells: generating and regenerating the brain. Neuron 80, 588–601.

Garg, H.G., Linhardt, R.J., and Hales, C.A. (2011). Chemistry and Biology of Heparin and Heparan Sulfate (Elsevier).

Gotz, J., and Ittner, L.M. (2008). Animal models of Alzheimer’s disease and frontotemporal dementia. Nat Rev Neurosci 9, 532–544.

Haass, C., and Selkoe, D.J. (2007). Soluble protein oligomers in neurodegeneration: lessons from the Alzheimer’s amyloid beta-peptide. Nat Rev Mol Cell Biol 8, 101–112.

Hardy, J., and Selkoe, D.J. (2002). The amyloid hypothesis of Alzheimer’s disease: progress and problems on the road to therapeutics. Science 297, 353–356.

Haughey, N.J., Liu, D., Nath, a., Borchard, a.C., and Mattson, M.P. (2002). Disruption of neurogenesis in the subventricular zone of adult mice, and in human cortical neuronal precursor cells in culture, by amyloid beta-peptide: implications for the pathogenesis of Alzheimer’s disease. Neuromolecular Med 1, 125–135.

Haycock, J.W. (2011). 3D cell culture: a review of current approaches and techniques. Methods Mol Biol 695, 1–15.

Heneka, M.T., Carson, M.J., El Khoury, J., Landreth, G.E., Brosseron, F., Feinstein, D. L., Jacobs, A.H., Wyss-Coray, T., Vitorica, J., Ransohoff, R.M., et al. (2015). Neuroinflammation in Alzheimer’s disease. The Lancet Neurology 14, 388–405.

Heo, C., Chang, K.-A., Choi, H.S., Kim, H.-S., Kim, S., Liew, H., Kim, J.a., Yu, E., Ma, J., and Suh, Y.-H. (2007). Effects of the monomeric, oligomeric, and fibrillar Aβ42 peptides on the proliferation and differentiation of adult neural stem cells from subventricular zone. Journal of Neurochemistry 102, 493–500.

Justice, B.A., Badr, N.A., and Felder, R.A. (2009). 3D cell culture opens new dimensions in cell-based assays. Drug Discov Today 14, 102–107.

Kienlen-Campard, P., Miolet, S., Tasiaux, B., and Octave, J.N. (2002). Intracellular amyloid-beta 1-42, but not extracellular soluble amyloid-beta peptides, induces neuronal apoptosis. J Biol Chem 277, 15666–15670.

Kim, J., Efe, J.A., Zhu, S., Talantova, M., Yuan, X., Wang, S., Lipton, S.A., Zhang, K., and Ding, S. (2011). Direct reprogramming of mouse fibroblasts to neural progenitors. Proc Natl Acad Sci U S A 108, 7838–7843.

Kim, K., Doi, A., Wen, B., Ng, K., Zhao, R., Cahan, P., Kim, J., Aryee, M.J., Ji, H., Ehrlich, L.I., et al. (2010). Epigenetic memory in induced pluripotent stem cells. Nature 467, 285–290.

Kizil, C., Iltzsche, A., Kuriakose, A., Bhattarai, P., Zhang, Y., and Brand, M. (2015). Efficient cargo delivery using a short cell-penetrating peptide in vertebrate brains. PLoS One 10, e0124073.

Koutsopoulos, S., and Zhang, S. (2013). Long-term three-dimensional neural tissue cultures in functionalized self-assembling peptide hydrogels, matrigel and collagen I. Acta Biomater 9, 5162–5169.

LaFerla, F.M., and Green, K.N. (2012). Animal models of Alzheimer disease. Cold Spring Harbor perspectives in medicine 2.

LaFerla, F.M., Green, K.N., and Oddo, S. (2007). Intracellular amyloid-beta in Alzheimer’s disease. Nat Rev Neurosci 8, 499–509.

Lau, L.W., Cua, R., Keough, M.B., Haylock-Jacobs, S., and Yong, V.W. (2013). Pathophysiology of the brain extracellular matrix: a new target for remyelination. Nat Rev Neurosci 14, 722–729.

Liao, Y., Smyth, G.K., and Shi, W. (2013). The Subread aligner: fast, accurate and scalable read mapping by seed-and-vote. Nucleic Acids Res 41, e108.

Liao, Y., Smyth, G.K., and Shi, W. (2014). featureCounts: an efficient general purpose program for assigning sequence reads to genomic features. Bioinformatics 30, 923–930.

Lilja, A.M., Malmsten, L., Röjdner, J., Voytenko, L., Verkhratsky, A., Ögren, S.O., Nordberg, A., and Marutle, A. (2015). Neural Stem Cell Transplant-Induced Effect on Neurogenesis and Cognition in Alzheimer Tg2576 Mice Is Inhibited by Concomitant Treatment with Amyloid-Lowering or Cholinergic α7 Nicotinic Receptor Drugs. Neural Plast 2015, 370432.

Lindvall, O., and Kokaia, Z. (2006). Stem cells for the treatment of neurological disorders. Nature 441, 1094–1096.

Love, M.I., Huber, W., and Anders, S. (2014). Moderated estimation of fold change and dispersion for RNA-seq data with DESeq2. Genome biology 15, 550.

Luo, W., and Brouwer, C. (2013). Pathview: an R/Bioconductor package for pathway-based data integration and visualization. Bioinformatics 29, 1830–1831.

Lutolf, M.P., Gilbert, P.M., and Blau, H.M. (2009). Designing materials to direct stem-cell fate. Nature 462, 433–441.

Maitz, M.F., Freudenberg, U., Tsurkan, M.V., Fischer, M., Beyrich, T., and Werner, C. (2013). Bio-responsive polymer hydrogels homeostatically regulate blood coagulation. Nat Commun 4, 2168.

Molyneaux, B.J., Arlotta, P., Menezes, J.R., and Macklis, J.D. (2007). Neuronal subtype specification in the cerebral cortex. Nat Rev Neurosci 8, 427–437.

Nalbantoglu, J., Tirado-Santiago, G., Lahsaini, A., Poirier, J., Goncalves, O., Verge, G., Momoli, F., Welner, S.A., Massicotte, G., Julien, J.P., et al. (1997). Impaired learning and LTP in mice expressing the carboxy terminus of the Alzheimer amyloid precursor protein. Nature 387, 500–505.

Ravi, M., Paramesh, V., Kaviya, S.R., Anuradha, E., and Solomon, F.D. (2015). 3D cell culture systems: advantages and applications. J Cell Physiol 230, 16–26.

Selkoe, D.J. (2002). Alzheimer’s disease is a synaptic failure. Science 298, 789–791.

Selkoe, D.J. (2003). Folding proteins in fatal ways. Nature 426, 900–904.

Smith, I., Silveirinha, V., Stein, J.L., de la Torre-Ubieta, L., Farrimond, J.A., Williamson, E.M., and Whalley, B.J. (2015). Human neural stem cell-derived cultures in three-dimensional substrates form spontaneously functional neuronal networks. J Tissue Eng Regen Med.

Tang-Schomer, M.D., White, J.D., Tien, L.W., Schmitt, L.I., Valentin, T.M., Graziano, D.J., Hopkins, A.M., Omenetto, F.G., Haydon, P.G., and Kaplan, D.L. (2014). Bioengineered functional brain-like cortical tissue. Proc Natl Acad Sci U S A 111, 13811–13816.

Tincer, G., Mashkaryan, V., Bhattarai, P., and Kizil, C. (2016). Neural stem/progenitor cells in Alzheimer’s disease. Yale J Biol Med 89, 23–35.

Tong, L.M., Fong, H., and Huang, Y. (2015). Stem cell therapy for Alzheimer’s disease and related disorders: current status and future perspectives. Experimental & molecular medicine 47, e151.

Tsurkan, M.V., Chwalek, K., Levental, K.R., Freudenberg, U., and Werner, C. (2011). Modular StarPEG-Heparin Gels with Bifunctional Peptide Linkers. Macromol Rapid Commun 31, 1529–1533.

Tsurkan, M.V., Chwalek, K., Prokoph, S., Zieris, A., Levental, K.R., Freudenberg, U., and Werner, C. (2013). Defined polymer-peptide conjugates to form cell-instructive starPEG-heparin matrices in situ. Adv Mater 25, 2606–2610.

Tsurkan, M.V., Levental, K.R., Freudenberg, U., and Werner, C. (2010). Enzymatically degradable heparin-polyethylene glycol gels with controlled mechanical properties. Chem Commun (Camb) 46, 1141–1143.

Venable, J.H., and Coggeshall, R. (1965). A Simplified Lead Citrate Stain for Use in Electron Microscopy. J Cell Biol 25, 407–408.

Verpelli, C., Carlessi, L., Bechi, G., Fusar Poli, E., Orellana, D., Heise, C., Franceschetti, S., Mantegazza, R., Mantegazza, M., Delia, D., et al. (2013). Comparative neuronal differentiation of self-renewing neural progenitor cell lines obtained from human induced pluripotent stem cells. Front Cell Neurosci 7, 175.

Wieduwild, R., Tsurkan, M., Chwalek, K., Murawala, P., Nowak, M., Freudenberg, U., Neinhuis, C., Werner, C., and Zhang, Y. (2013). Minimal peptide motif for non-covalent peptide-heparin hydrogels. J Am Chem Soc 135, 2919–2922.

Wu, T.D., Reeder, J., Lawrence, M., Becker, G., and Brauer, M.J. (2016). GMAP and GSNAP for Genomic Sequence Alignment: Enhancements to Speed, Accuracy, and Functionality. Methods Mol Biol 1418, 283–334.

Wyss-Coray, T. (2016). Ageing, neurodegeneration and brain rejuvenation. Nature 539, 180–186.

Xia, N., Zhang, P., Fang, F., Wang, Z., Rothstein, M., Angulo, B., Chiang, R., Taylor, J., and Reijo Pera, R.A. (2016). Transcriptional comparison of human induced and primary midbrain dopaminergic neurons. Sci Rep 6, 20270.

Young, M.D., Wakefield, M.J., Smyth, G.K., and Oshlack, A. (2010). Gene ontology analysis for RNA-seq: accounting for selection bias. Genome biology 11, R14.

Yu, G., Wang, L.G., Han, Y., and He, Q.Y. (2012). clusterProfiler: an R package for comparing biological themes among gene clusters. OMICS 16, 284–287.

Zhang, D., Pekkanen-Mattila, M., Shahsavani, M., Falk, A., Teixeira, A.I., and Herland, A. (2014). A 3D Alzheimer’s disease culture model and the induction of P21-activated kinase mediated sensing in iPSC derived neurons. Biomaterials 35, 1420–1428.

Zhang, Y., Fussel, S., Reimer, U., Schutkowski, M., and Fischer, G. (2002). Substrate-based design of reversible Pin1 inhibitors. Biochemistry 41, 11868–11877.

